# High-pass filtering artifacts in multivariate classification of neural time series data

**DOI:** 10.1101/530220

**Authors:** Joram van Driel, Christian N.L. Olivers, Johannes J. Fahrenfort

**Affiliations:** Institute for Brain and Behaviour Amsterdam, Department of Experimental and Applied Psychology - Cognitive Psychology, Faculty of Behavioural and Movement Sciences, Vrije Universiteit Amsterdam

**Keywords:** EEG, MEG, high-pass filtering, preprocessing, multivariate pattern classification, decoding

## Abstract

0.

**Background:** Traditionally, EEG/MEG data are high-pass filtered and baseline-corrected to remove slow drifts. Minor deleterious effects of high-pass filtering in traditional time-series analysis have been well-documented, including temporal displacements. However, its effects on time-resolved multivariate pattern classification analyses (MVPA) are largely unknown.

**New Method:** To prevent potential displacement effects, we extend an alternative method of removing slow drift noise – robust detrending – with a procedure in which we mask out all cortical events from each trial. We refer to this method as *trial-masked robust detrending*.

**Results:** In both real and simulated EEG data of a working memory experiment, we show that both high-pass filtering and standard robust detrending create artifacts that result in the displacement of multivariate patterns into activity silent periods, particularly apparent in temporal generalization analyses, and especially in combination with baseline correction. We show that trial-masked robust detrending is free from such displacements.

**Comparison with Existing Method(s):** Temporal displacement may emerge even with modest filter cut-off settings such as 0.05 Hz, and even in regular robust detrending. However, trial-masked robust detrending results in artifact-free decoding without displacements. Baseline correction may unwittingly obfuscate spurious decoding effects and displace them to the rest of the trial.

**Conclusions:** Decoding analyses benefits from trial-masked robust detrending, without the unwanted side effects introduced by filtering or regular robust detrending. However, for sufficiently clean data sets and sufficiently strong signals, no filtering or detrending at all may work adequately. Implications for other types of data are discussed, followed by a number of recommendations.

## 1. Introduction

Recent years have seen an upsurge in the application of time-resolved multivariate pattern classification analyses (MVPA) – also referred to as *decoding* – to electro- and magnetoencephalographic data (EEG/MEG; see Table 1 for an extensive list of references). MVPA allows researchers to uncover the active sensory and mnemonic representations underlying cognitive processes as wide-ranging as perception, attention, categorization, language, working memory, and long-term memory. Many researchers therefore now prefer the information-rich multivariate approach over traditional univariate event-related potential (ERP) or event-related field (ERF) analyses based on signals averaged over epochs. Moreover, a number of toolboxes have recently emerged to facilitate these types of analyses (e.g. Bode, Feuerriegel, Bennett, & Alday, 2018; Fahrenfort, van Driel, van Gaal, & Olivers, 2018; Hanke et al., 2009; Meyers, 2013; Oosterhof, Connolly, & Haxby, 2016; Treder, 2020).

**Table 1.** A non-exhaustive list of EEG and MEG studies that have used MVPA decoding techniques after applying different levels of filtering. High-pass and low-pass cut-off values are provided. Note this table is only intended to illustrate the wide-ranging use of high-pass filters in EEG/MEG, and not to suggest that anything is necessarily wrong with these studies. For example, different studies may use different filter types: online (causal) or offline (either causal or acausal), Finite Impulse Response (FIR) or Infinite Impulse Response (IIR), different filter lengths and so forth, and each of these filter types may have different effects on the data that do not necessarily have to be problematic in the scientific context in which they are applied.

| *Publication* | *EEG* | *MEG* | *High pass cut-off (Hz)* | *Low pass cut-off (Hz)* |
| --- | --- | --- | --- | --- |
| (Alizadeh, Jamalabadi, Schonauer, Leibold, & Gais, 2017) | • |  | 0.1 | 40 |
| (Aukstulewicz, Myers, Schnupp, & Nobre, 2019) | • | • | 0.1 | 200 |
| (Bae & Luck, 2018) | • |  | 0.1 | 80 |
| (Bae & Luck, 2019) | • |  | 0.1 | 80 |
| (Barragan-Jason, Cauchoix, & Barbeau, 2015) | • |  | 0.1 | 40 |
| (Blom et al., 2020) | • |  | - | - |
| (Boettcher, Stokes, Nobre, & van Ede, 2020) | • |  | 0.1 | - |
| (Borst, Ghuman, & Anderson, 2016) |  | • | 0.5 | 50 |
| (Borst, Schneider, Walsh, & Anderson, 2013) | • |  | 0.5 | 30 |
| (Brandman, Avancini, Leticevscaia, & Peelen, 2020) |  | • | 0.1 | 330 |
| (Brandmeyer, Farquhar, McQueen, & Desain, 2013) | • |  | 1 | 25 |
| (Carlson, Hogendoorn, Kanai, Mesik, & Turret, 2011) |  | • | - | - |
| (Carlson, Tovar, Alink, & Kriegeskorte, 2013) |  | • | 0.1 | 200 |
| (Cauchoix, Barragan-Jason, Serre, & Barbeau, 2014) |  |  | 0.1 | 40 |
| (Chan, Halgren, Marinkovic, & Cash, 2011) | • | • | 0.1 | 200 |
| (Cichy & Pantazis, 2017) | • | • | 0.03 | 300 |
| (Cichy, Pantazis, & Oliva, 2014) |  | • | 0.03 | 330 |
| (Cichy, Ramirez, & Pantazis, 2015) |  | • | 0.03 | 330 |
| (Clarke, Devereux, Randall, & Tyler, 2015) |  | • | 0.03 | 40 |
| (Correia, Jansma, Hausfeld, Kikkert, & Bonte, 2015) | • |  | 0.1 | 100 |
| (Dash, Ferrari, & Wang, 2020) |  | • | - | 250 |
| (Dijkstra, Mostert, Lange, Bosch, & van Gerven, 2018) |  | • | - | - |
| (Fahrenfort, Grubert, Olivers, & Eimer, 2017) | • |  | 0.1 | - |
| (Fahrenfort, van Leeuwen, et al., 2017) | • |  | 0.1 | - |
| (Giari, Leonardelli, Tao, Machado, & Fairhall, 2020) |  | • | 0.8 | 80 |
| (Gwilliams & King, 2020) |  | • | 0.5 | 40 |
| (Hebart, Bankson, Harel, Baker, & Cichy, 2018) |  | • | 0.1 | 300 |
| (Herrmann, Maess, Kalberlah, Haynes, & Friederici, 2012) |  | • | 2 | 10 |
| (Hogendoorn & Burkitt, 2018) | • |  | - | - |
| (Hogendoorn, Verstraten, & Cavanagh, 2015) | • |  | - | - |
| (Hubbard, Kikumoto, & Mayr, 2019) | • |  | 0.01 | 80 |
| (Isik, Meyers, Leibo, & Poggio, 2014) |  | • | 2 | 100 |
| (Jach, Feuerriegel, & Smillie, 2020) | • |  | 0.5 | 30 |
| (Jensen, Hyder, & Shtyrov, 2019) |  | • | 1 | 95 |
| (Kaiser, Azzalini, & Peelen, 2016) |  | • | 1 | 330 |
| (Kaiser, Oosterhof, & Peelen, 2016) |  | • | 1 | 300 |
| (Kaiser, Moeskops, & Cichy, 2018) | • |  | 0.5 | 70 |
| (King, Pescetelli, & Dehaene, 2016) |  | • | 0.1 | 30 |
| (Kok, Mostert, & de Lange, 2017) |  | • | - | 40 |
| (LaRocque, Lewis-Peacock, Drysdale, Oberauer, & Postle, 2013) | • |  | 1 | 55 |
| (Liang, Oba, & Ishii, 2019) | • |  | 1 | 45 |
| (Ling, Lee, Armstrong, & Nestor, 2019) | • |  | 0.1 | 40 |
| (Mai, Grootswagers, & Carlson, 2019) |  | • | 0.03 | 200 |
| (Mares, Ewing, Farran, Smith, & Smith, 2020) | • |  | 0.1 | 40 |
| (Marti & Dehaene, 2017) |  | • | 0.1 | 30 |
| (Marti, King, & Dehaene, 2015) |  | • | 0.1 | 330 |
| (Mohsenzadeh, Qin, Cichy, & Pantazis, 2018) |  | • | 0.03 | 330 |
| (Mostert, Kok, & de Lange, 2015) |  | • | - | 30 |
| (Munneke, Fahrenfort, Sutterer, Theeuwes, & Awh, 2020) | • |  | 0.01 | 80 |
| (Myers et al., 2015) | • | • | 0.03 | 300 |
| (Nemrodov, Niemeier, Mok, & Nestor, 2016) | • |  | 0.1 | 40 |
| (Nemrodov, Niemeier, Patel, & Nestor, 2018) | • |  | 0.1 | 40 |
| (Noah et al., 2020) | • |  | - | 40 |
| (Ort, Fahrenfort, Ten Cate, Eimer, & Olivers, 2019) | • |  | - | - |
| (Peters, Bledowski, Rieder, & Kaiser, 2016) |  | • | 0.1 | 150 |
| (Quax, Dijkstra, van Staveren, Bosch, & van Gerven, 2019) |  | • | - | - |
| (Rose et al., 2016) | • |  | 1 | 60 |
| (Simanova, van Gerven, Oostenveld, & Hagoort, 2010) | • |  | 1 | 30 |
| (Sudre et al., 2012) |  | • | 0.1 | 50 |
| (Takacs, Mückschel, Roessner, & Beste, 2020) | • |  | 0.5 | 40 |
| (Tankelevitch, Spaak, Rushworth, & Stokes, 2020) |  | • | 0.1 | 40 |
| (Teichmann, Grootswagers, Carlson, & Rich, 2019) |  | • | 0.03 | 200 |
| (Treder, 2020) | • | • | 0.1 | 100 |
| (Trubutschek et al., 2017) |  | • | 0.1 | 330 |
| (Turner, Johnston, de Boer, Morawetz, & Bode, 2017) | • |  | 0.1 | 70 |
| (Wardle, Kriegeskorte, Grootswagers, Khaligh-Razavi, & Carlson, 2016) |  | • | 0.1 | 200 |
| (Wolff et al., 2015) | • |  | 0.1 | 40 |
| (Wolff, Jochim, Akyurek, & Stokes, 2017) | • |  | 0.1 | 40 |

However, as the field is making the transition from univariate to multivariate approaches, some of the standard data processing procedures remain, raising the question whether these procedures are actually optimal, or perhaps even harmful, for decoding. One of the most common processing steps is *high-pass filtering*. Given the slow drifts in especially EEG data (less so in MEG data), high-pass filtering has become a crucial component in extracting ERPs and improving signal-to-noise. However, it is well known that high-pass filtering can lead to artifacts. Specifically, too high cut-off values (typically 0.1 Hz or more) may cause the signal enhancement to result in spurious local ringing effects^1^ around the event-related responses – artifacts which may be misinterpreted as real components in the event-related signal (Acunzo, MacKenzie, & van Rossum, 2012; Kappenman & Luck, 2010; Luck, 2005; Tanner, Morgan-Short, & Luck, 2015; Tanner, Norton, Morgan-Short, & Luck, 2016; Widmann, Schroger, & Maess, 2015). Nevertheless, high-pass filtering is generally still considered a crucial step for extracting meaningful ERPs (for which drift correction is necessary), and therefore continues to be part of the recommendations with regards to EEG data preprocessing (with appropriate cut-off values, e.g. Maess, Schroger, & Widmann, 2016; Tanner et al., 2016; Widmann & Schroger, 2012; Widmann et al., 2015).

Perhaps less well known is that, depending on the specific cut-off value and frequency of the ERP, high-pass filtering may also lead to quite diffuse, but still spurious, activity differences both well before and well after the event-related response (Tanner et al., 2016). Even with modest cut-off settings, these slower components may emerge as subtle overall baseline shifts. A not uncommon step for ERP researchers is to correct for these shifts (whether apparent or real), thus potentially obscuring any artifacts. Thus, although sufficiently powered ERP studies could still show such artifacts, subtle baseline differences are often thought to be remedied by ensuing baseline corrections in ERP analyses (though see Tanner et al., 2016). However, multivariate analyses may be more sensitive to spuriously transposed information present in the topographical landscape. So far, little is known about the effects of high-pass filtering on multivariate pattern classification, and to what extent it leads to artifacts in decoding.

The potential for spurious temporal displacement of information is particularly worrisome when testing hypotheses on neural activity in the absence of stimulation, for example in the field of working memory. Indeed, after extensively analyzing one of our own EEG-based working memory experiments, we had to conclude that the above-chance decoding of the memoranda during the blank delay period was at least partly caused by the (modest) high-pass filter applied during preprocessing. As Table 1 shows, we have not been the only ones applying high-pass filtering prior to MVPA, as filtering has remained part of the pre-processing pipeline in a wide range of studies. Moreover, the same table also shows a wide range of cut-off values used when high-pass filtering is applied, from as low as 0.03 Hz to as high as 2 Hz, with 0.1 Hz being the most typical^2^. We thus decided to conduct a systematic exploration of high-pass filtering-related artifacts in MVPA, the results of which are presented here. First, we show how high-pass filtering led to clear signs of spurious decoding in one of our own EEG experiments, which involved a working memory task illustrated in Figure 1. The task contained an initial presentation of a cue, a blank delay period during which the cue had to be retained, and a test stimulus in which observers searched for the cued object. To uncover the cause of the artifacts, and because empirical data does not come with a ground truth, we subsequently chose to create a simulated data set that allowed us to assess how decoding of filtered signals compares against decoding a known raw signal.

**Figure 1.**
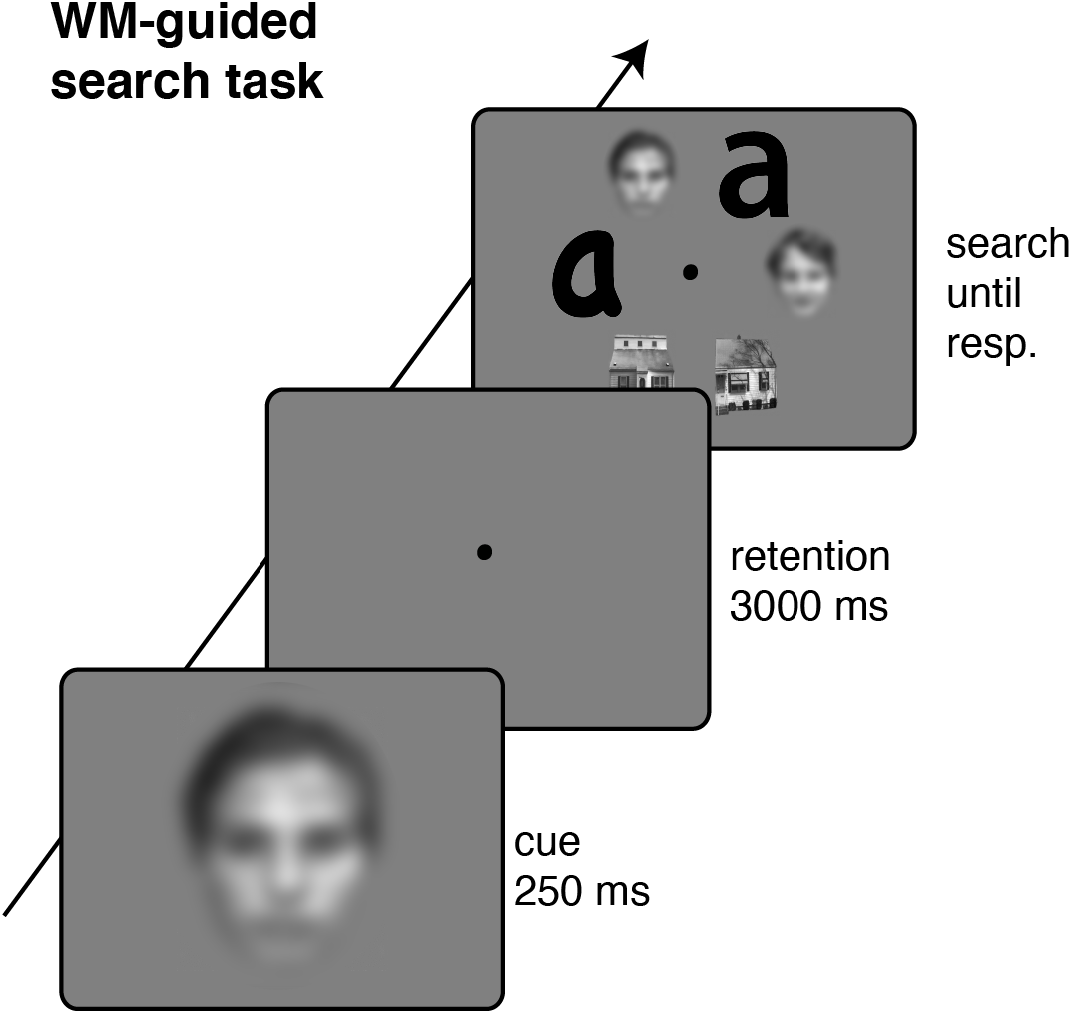
Example trial for the empirical experiment. Observers remembered a cued house, face or letter target for a subsequent visual search task presented after a 3000 ms blank retention interval. Observers then indicated with one of two button presses whether the memorized target was present or absent in a visual search display which also contained a number of nontarget objects. The faces in the search display were replaced with blurred versions in this illustration to make them unidentifiable for reasons of privacy

In addition to testing the effects of high-pass filtering, we tested two alternative methods to remove slow drifts: *robust detrending*, and an extension of this method which we refer to as *trial-masked robust detrending*. In detrending, an n^th^ order polynomial is fitted to the data and subsequently subtracted from the data, thereby removing slow-fluctuating drifts. Because such fits can be sensitive to artifactual deviations (glitches) from the slow trend, Cheveigné and Arzounian (2018) recently introduced an improved method called *robust detrending* in which they employ an iterative weighting procedure to mask outliers from the data to which the polynomial is fitted. Although (robust) detrending is preferable over high-pass filtering, this method can still affect the data in at least two undesirable ways: (1) A slow polynomial may shift upwards or downwards by fitting to the peak of the ERP, thereby slightly moving activity from the peak of the ERP to the temporal window over which the shift occurred when the polynomial is subtracted and/or (2) a polynomial may fit to a long-duration low amplitude ERP such that (some of) the ERP itself is subtracted out when the polynomial is subtracted out.

Therefore, we extend the robust detrending method in the current manuscript by not only masking out glitches that are determined in a data-driven way, but by actively masking out all parts of the data that might contain relevant cortical events (such as ERPs). We call this method *trial-masked robust detrending*, which should in principle preclude any kind of influence of experimentally relevant events on the detrending procedure. We show that both high-pass filtering and standard robust detrending can result in multiple-comparison FDR (False Discovery Rate) corrected spurious decoding in time intervals where no above chance decoding should be present (such as the baseline window). These effects are particularly strong in temporal generalization matrices. In contrast, trial-masked robust detrending does not result in such spurious decoding effects, while still realizing modest improvements in decoding performance. Furthermore, we show that spurious effects in the baseline window may be transported to the rest of the trial through baseline-correction, thus unwittingly obfuscating spurious effects that were introduced by the pre-processing method.

## 2. Methods

For both the empirical and the simulated data set, stimuli, data, code and analyses scripts are available from the Open Science Framework, at https://osf.io/t9rkz/.

### 2.1. Empirical data

We report data from an experiment that is illustrated in Figure 1. On every trial, observers were presented with a face, house, or letter (the cue), which they had to remember for a visual search task presented 3 seconds later. The task was to determine the presence or absence of the cued target. The experiment included other conditions, but to simplify matters here we report on the condition that best serves the current purpose.

#### 2.1.1. Participants

Twenty-five students from the Vrije Universiteit Amsterdam participated for course credits or monetary payment (€9 per hour). All subjects reported normal or corrected to normal vision. The protocol complied with ethical guidelines as approved by the Scientific and Ethical Review Committee of the Faculty of Behavioural and Movement Sciences, and with the Declaration of Helsinki. Data of two subjects were removed from further analyses, one due to excessive high frequency noise reflecting muscle artifacts, and another due to a very strong but poorly understood artifact in the ERPs that is most likely due to equipment failure.

#### 2.1.2. Stimuli and task

Subjects were asked to memorize at each trial a briefly presented picture (250 ms), which could be of the category face, house or letter (width: ~4° visual angle; height: ~5°). After a retention interval of 3 seconds (with only a white dot at the center of the screen as fixation point), a search display appeared, consisting of six pictures (two exemplars of each category; ~2.5° in size) randomly arranged along a hexagon array (radius of 4.5°; three pictures per hemifield; white fixation dot remained at the center of the screen). Subjects were asked to indicate whether the target picture they memorized at the start of the trial was present (left index finger) or absent (right index finger) by pressing a button on a button box connected to the EEG acquisition computer via a parallel port. Probability of target present/absent was 50%. The search array disappeared upon the subject’s response (which changed the color of the fixation dot to black for 500 ms), or when 5 seconds had passed (after which the warning “respond faster!” appeared at the center of the screen for 500 ms). The inter-trial interval was set to 1 second ± 500 ms jitter. Low-level image properties of face and house pictures were controlled with the SHINE toolbox (Willenbockel et al., 2010). Subjects performed a short practice block of 12 trials with feedback on accuracy (words “correct!” and “wrong…” presented centrally for 500 ms), after which EEG recording started for 252 trials (84 per picture category), without feedback (except for slow responses). Prior to participating, subjects signed an informed consent form. Each unique picture within a category was only presented once as target, while it could be used more than once as distractors within the search arrays. Furthermore, when the target was a face, the two face stimuli in the search display were of same gender, encouraging subjects to memorize facial features rather than category. We randomly selected face stimuli out of 100 face pictures (from Endl et al., 1998, 50 male, 50 female). Similarly, when the target was a letter, the two letter stimuli in the search display where of same identity and capitalization, encouraging subjects to memorize the specific font. House stimuli were randomly sampled from 100 exemplars of pictures used in Egner, Monti, and Summerfield (2010).

#### 2.1.3. Data acquisition

EEG data from 64 Biosemi ActiveTwo (biosemi.com) electrodes placed according to the 10-5 system (a high-resolution electrode placement standard derived from the international 10-20 system) were acquired at 512 Hz sampling rate. The ActiveTwo system is DC-coupled, and thus has no online (hardware) high-pass filter. On such DC-coupled systems, drifts are common due to non-brain artifacts such as sweating. Further, the data was down-sampled offline to 128 Hz and re-referenced to the average of signals recorded from both earlobes. Error trials, trials without a response, or with responses slower than 3 seconds were not included in the analyses. Continuous, raw data was first inspected for malfunctioning electrodes, which were interpolated after the below preprocessing steps. We did not perform any oculomotor artifact correction.

### 2.2. Simulated data

We describe the creation of an artificial dataset for a task with a highly similar data structure. As a basis for the simulated data, we took the continuous EEG data structure from a representative subject and replaced the continuous EEG of that subject with simulated EEG, with ERPs injected at the same points in time where ERPs occurred in the original dataset. Each of the simulated ERPs involved the simulated presentation of a to-be-remembered stimulus, a retention phase, an activity silent period and a test stimulus (described below in section 2.2.1 and 2.2.2 below). The continuous signal also contained simulated continuous pink noise (described below in section 2.2.3 below).

#### 2.2.1. Creating class-specific topographical patterns

Figure 2A illustrates the creation of the underlying spatial pattern of evoked responses. The features fed into a linear discriminant classifier are typically activity values at a given time point (or averaged over a time window) for each of N electrodes that cover the scalp or part thereof. Here we simulated activity of 64 electrodes. From these we selected a fixed set of electrodes to represent one stimulus class, and another, partially overlapping set of electrodes to represent another stimulus class. Stimulus-related class-specific activity was thus associated with different multivariate spatial patterns, such that multivariate classification trained and tested on the channel features over time would be able to reproduce the stimulus-related activity.

**Figure 2.**
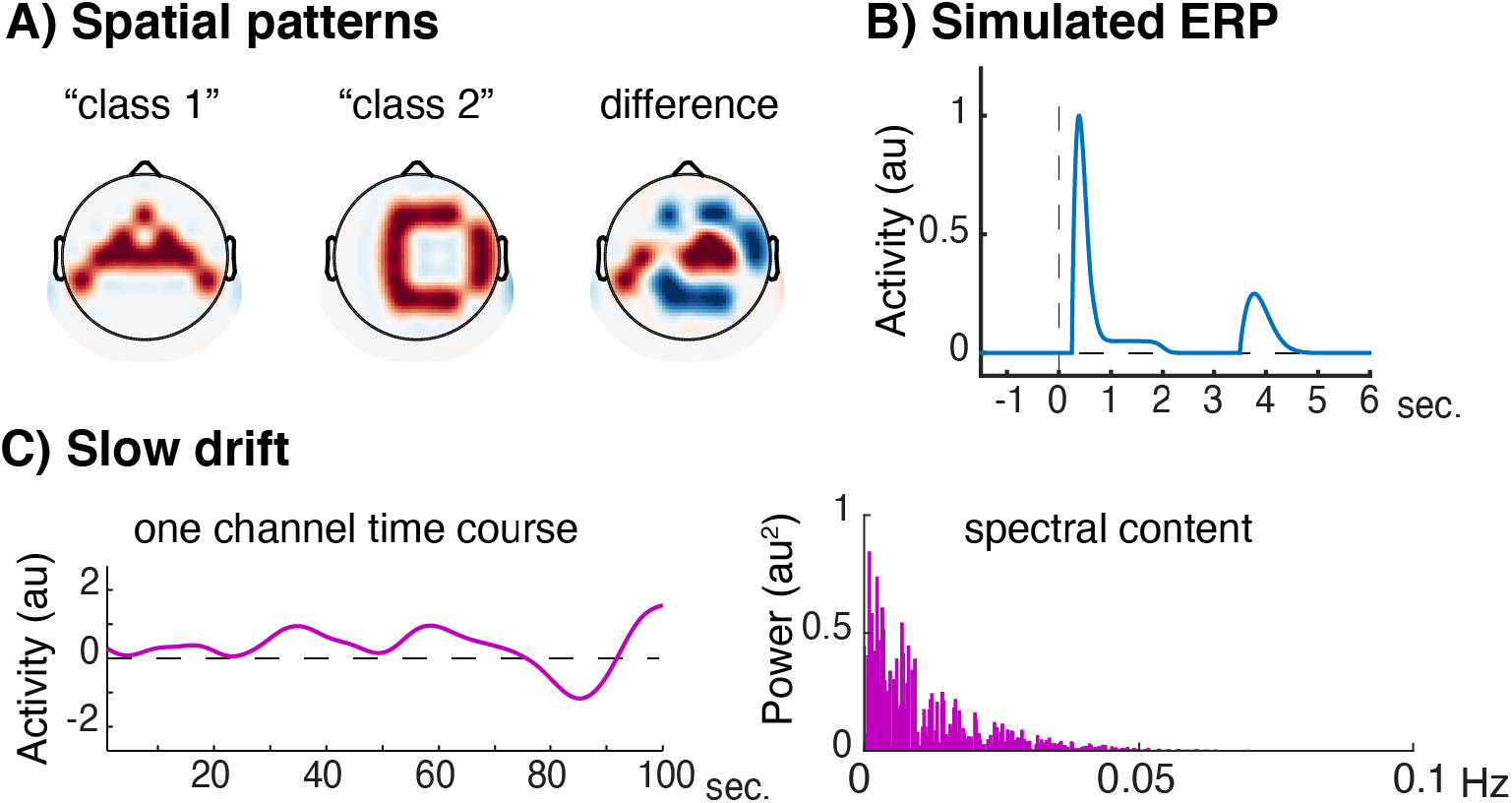
Creation of simulated data. A) Two different electrode topographies representing the two stimulus classes, plus their difference. Red as positive, blue as negative. B) The underlying simulated ERP time series as injected into each electrode of the topographical patterns. C) Example time course of 1/f pink noise slow drift as was added to the data (left panel), and its spectral content (right panel).

To simulate stimulus-related activity assigned to these sets of electrodes, we first created an event-related potential (ERP, shown in Figure 2B) that mimicked four phases of a working memory task: (1) encoding the stimulus into the visual system, (2) actively retaining a representation of the stimulus in working memory, (3) an activity silent period, and (4) a search phase in which subjects attempt to determine the presence of the target stimulus upon the presentation of a probe. A “trial” consisted of an array of data containing the entire ERP, and lasted from −1.5 seconds to 6 seconds surrounding the “event” (what would be the onset of the to-be-encoded stimulus). The four phases of the ERP were modeled as follows: the encoding response started at *t* = 0.25 seconds using a Weibull function with a steep rising slope peaking at amplitude 1 a.u. and a shallower falling slope dropping to baseline at *t* = 1 seconds. The active retention phase (partially overlapping with the encoding phase) started at *t* = 0.5 seconds, modeled by a shallower logistic curve containing a plateau with an amplitude of 0.05 a.u. continuing for about 1 second, and dropping back to baseline around *t* = 2.5 seconds. The silent retention period with activity at baseline lasted from *t* = 2.5 seconds until 3.5 seconds. The search phase (memory display) started at *t* = 3.5 seconds, modeled by a similar Weibull function as for encoding but with a lower amplitude of 0.25 a.u., dropping back to baseline at *t* = 5 seconds, and staying at baseline for the remainder of the trial. Note that the silent period continued for an extended period of time (1 second) to determine whether decoding would occur in a time period where no information was present in the original data.

#### 2.2.2. Decision boundary

We created one class of patterns by injecting (adding) the ERP into each of the electrodes in one of the two sets described above for 112 trials, and another class by injecting the ERP to each of the electrodes in the other set for 112 trials, thus creating two different spatiotemporal landscapes of activity. The order of the trials was randomized before injection. With such two highly different patterns, a classifier would produce perfect classification performance. To avoid such a ceiling effect, we took three measures (1) we compromised the distance of the classes to the decision boundary by warping the classes toward each other and (2) we varied the amplitude of the ERP for different phases in the trial (see previous section) and (3) we injected pink noise into the data (see next section). To compromise the distance to the decision boundary, we first created a “decision boundary space” by warping one spatial pattern into the other pattern in 80 linearly spaced transitions. Warping resulted in non-overlapping channels now showing a *relatively* stronger ERP for one stimulus class than the other (where this was binary prior to warping). Another way of describing the effect of this warping procedure is that the multivariate patterns of the two stimulus classes become more similar, thereby moving them closer to the decision boundary of a multivariate classifier. We used intermediately divergent patterns for the two classes (transition 32 vs transition 49, where the most divergent patterns would be transition 1 vs 80 and the least divergent patterns would be 40 vs 41). The same patterns were used throughout the trial (to allow for temporal generalization), while the separability of the patterns in any given phase of the trial was determined by the signal to noise ratio in that phase (see next section). The warping parameter was not varied, so that the same distance to the decision boundary was used throughout the simulated experiment. Note that these simulated ERPs and spatial distributions were purely meant to illustrate decoding under separability, and not intended as an exact model of brain mechanisms of working memory. Nevertheless, the classification result yielded a pattern that can be observed in data from other labs (e.g. Myers et al., 2015; Wolff, Ding, Myers, & Stokes, 2015), as well as in our own data (see Figure 4): A transient sweep of high decoding performance during encoding, lower sustained performance during retention, an activity silent period, and high (yet reduced) performance during recall/search.

**Figure 3.**
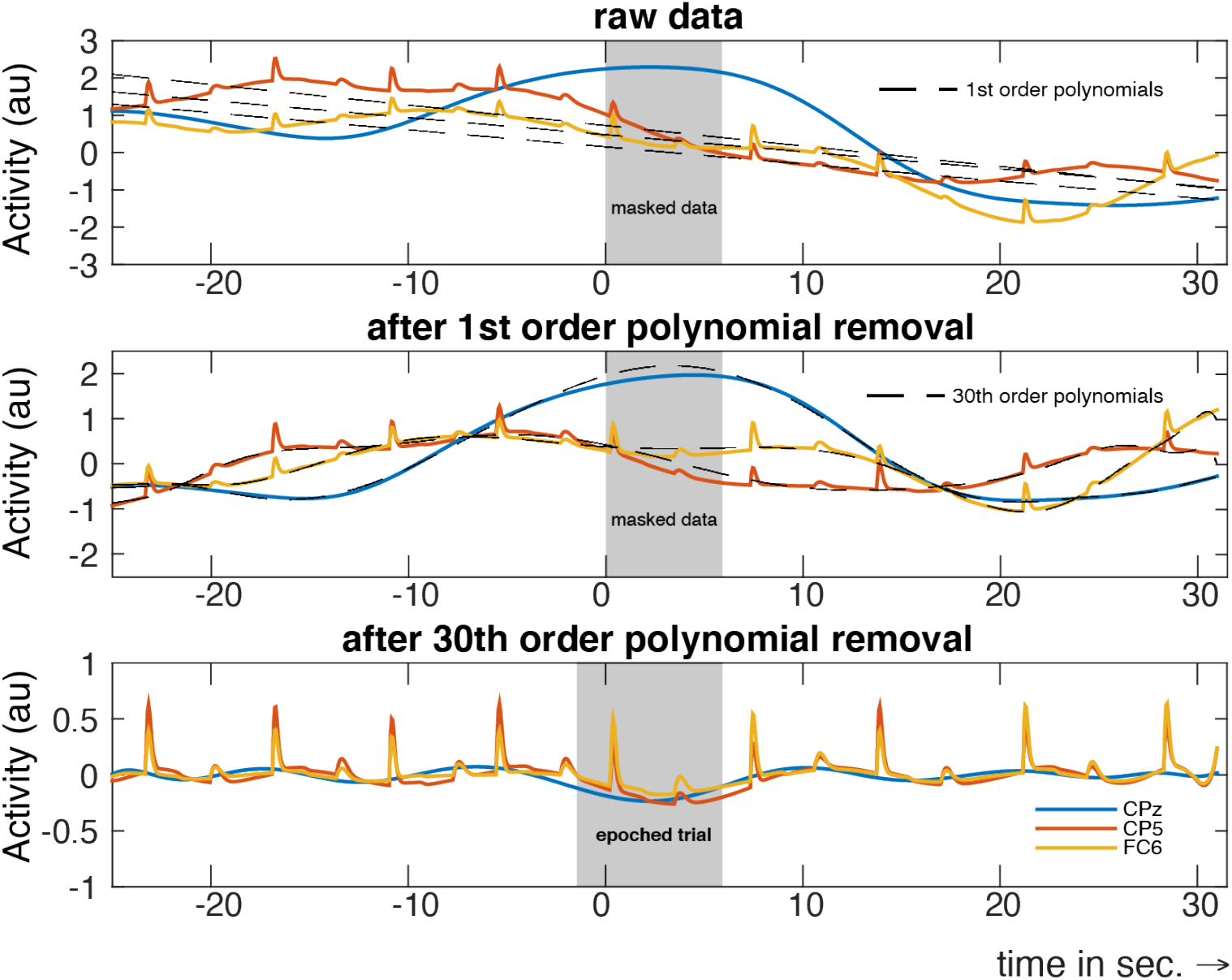
Procedure for removing low-frequency drift noise with current trial-masked robust detrending. This figure shows the detrending procedure on simulated data of an illustrative trial of some illustrative electrodes in an illustrative subject. Top panel: raw data for three electrodes: CPz (no ERP), CP5 and FC6 (both of which contain an ERP). Also shown as dotted lines are the polynomial fits on the raw data from which the trial events were masked out (grey panel in the background). Middle panel: data after removing 1^st^ order polynomials fits from the top panel. Also shown are the 30^th^ order polynomial fits on these data, from which the trial events were masked out (grey panel in the background). Bottom: data after removing the 30^th^ order polynomial fits. Finally, the middle trial is segmented out for further analysis. Note that the epoched trial is slightly wider than the mask that was used during detrending because it also includes a 1.5 second pre-stimulus period. Further, depending on the length of the intertrial interval one may choose to mask out only the data that are contained in the currently epoched trial (as was done in the analyses presented in this paper), or also mask out neighboring trials (which may also work well, but was not done here or in the analyses that were presented in this manuscript). Finally, the robust detrending algorithm will also iteratively mask out additional sharp transients from the data that otherwise disturb smooth fits, see main text as well as (de Cheveigne & Arzounian, 2018) for details. A similar figure is produced when using the detrending function included in the ADAM toolbox, an illustration of which can be found in Supplementary Figure S1.

**Figure 4.**
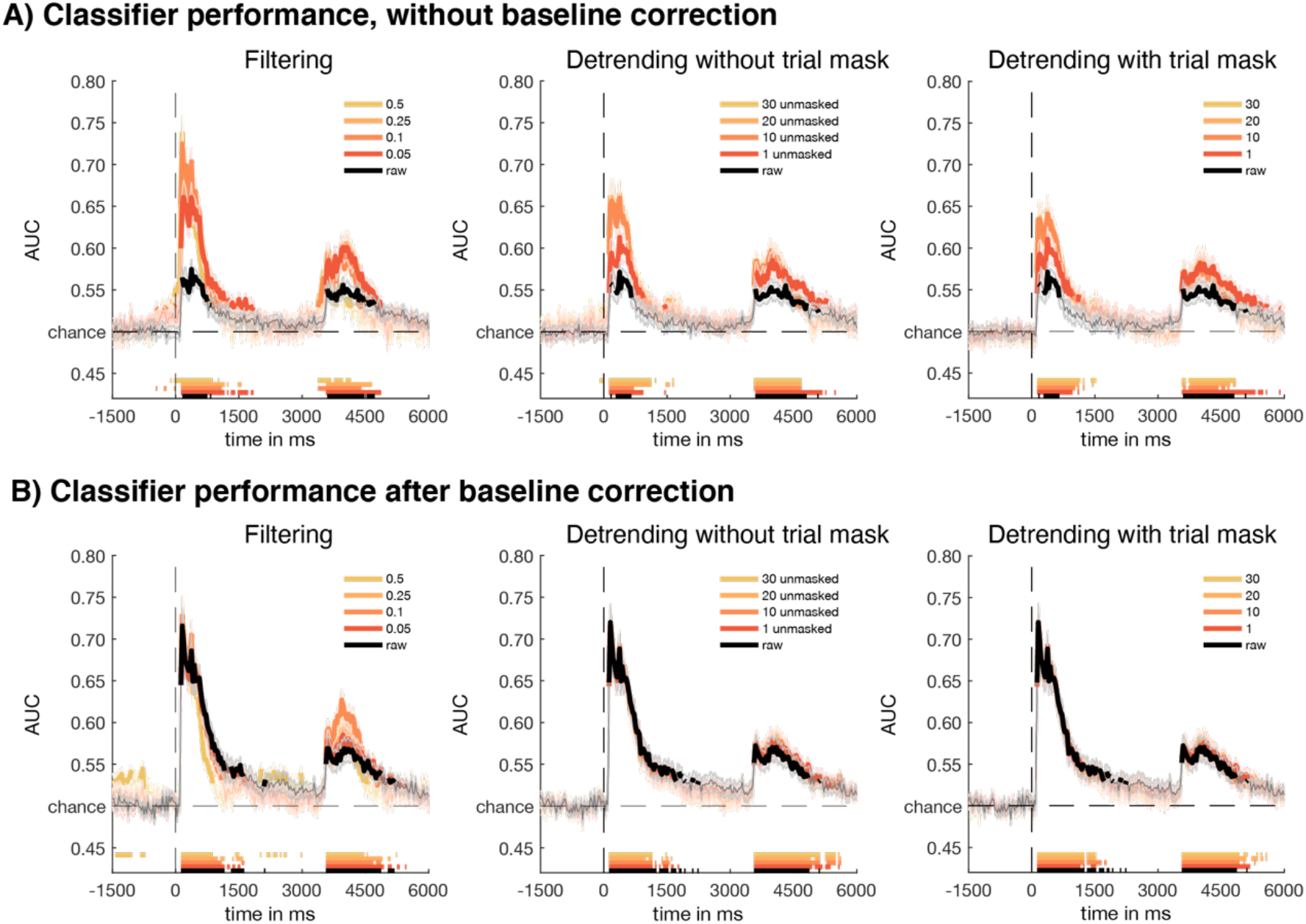
Results for empirical data from a working memory guided search task with and without prior baseline correction. A) Decoding performance (AUC) without baseline correction at each time point for different high-pass filter cut-off frequencies (left panel), regular robust detrending (middle panel) and trial-masked robust detrending (right panel). Different pre-processing parameters (high-pass filter cutoff / polynomial order) are indicated by shades of orange, while raw data is shown in black. All thick colored lines denote reliable difference from chance (p<0.05) after FDR correction (q=0.05). B) The same, but now after a baselinecorrection on a window of −200 to 0 milliseconds. Note reliable above-chance decoding at the onset of the trial in the high-pass filtered data, especially after a cut-off of 0.5 Hz. Although the artifacts may seem minor in the non-baselined data, baseline-correction itself may cause spurious effects to be transposed to the rest of the trial due to the subtraction logic of baseline correction (i.e. subtracting the average activity in an electrode during the baseline from the activity in the rest of the trial in fact sets the pattern of activity at baseline to zero and displaces the inverse of the original pattern that was present at baseline to the rest of the trial). Note that the baseline window in this example runs from −200 to 0 milliseconds, thus the above chance decoding performance for the 0.5 Hz cut-off at the beginning of the trial is plausibly the result of such a displacement. Together, this may also lead one to question whether the observed above-chance decoding during the retention period for the cut-off of 0.5 Hz is real. Moreover, the effects of such artifactual displacements on temporal generalization may be even worse (see Figure 5 and 6).

#### 2.2.3. Adding low-frequency pink drifts as noise

As high-pass filtering and robust detrending are used to remove low-frequency non-stationary drifts, the next step was to generate simulated data that contained such activity. To this end, we created a time series of pink noise (1/f, power spectral density is inversely related to frequency) for each electrode separately. Specifically, we first created Fourier coefficients of random amplitudes (drawn from a uniform distribution between 0 and 1) with an exponential decay over frequency, and multiplied these by random phase angles (drawn from a uniform circular distribution). The real part of the inverse fast Fourier transform of this simulated power spectrum produced “continuous data” of low frequency noise. The exponential decay function of the power spectrum had the form exp(−(0:nSamples-1)/rolloff), where nSamples is the number of Fourier coefficients (i.e. equal to the number of samples / sine waves used to generate the simulated signal), and rolloff determined the steepness of the decline. In our simulation, nSamples was 539648 and rolloff was 100. This decrease resulted in an attenuation of power of −3.65 dB when going from 0.0001 Hz to 0.1 Hz, such that power values at 0.1 Hz approximated zero (Figure 2D shows an illustrative 100 second snippet of the noise drift time series of one channel, and the corresponding power spectrum that was used to generate it).

To make the data structure of the simulated data comparable to data structure of the empirical data, we used the continuous data file of a representative subject and replaced its real data with simulated pink noise. The total length of this time series was therefore equal to the total length of the experiment of that subject (containing 539648 samples) utilizing the same sampling frequency (128 Hz), and thus resulting in a total of around 70 minutes of simulated data (539648 samples /128 Hz / 60 seconds = ~ 70 minutes). Next, we randomized the order of the events in the accompanying event structure, and added ERPs of the two classes where real trial events would have otherwise occurred in the empirical experiment, so that the inter-trial event structure in the simulated data was the same as in the empirical data.

Because it is impossible to recreate the exact signal to noise ratio of the real experiment due to mixing of signal and noise therein, we fixed the signal to noise ratio in any given electrode of any given participant by scaling the maximum amplitude of the simulated noise by a factor of 2.5 a.u. (compared to a maximum amplitude of 1 a.u. in the ERP). Further, because different phases of a trial had different ERP amplitudes associated with it (see previous section), the signal to noise ratio varied throughout the trial from −7.96 dB for the encoding phase, −33.98 dB for the retention phase and −20 dB for the search phase (under the assumption of a weight of 1 for that electrode). This variation makes it possible to inspect the effect of the different pre-processing procedures on decoding under different ratios of signal to noise. Finally, to able to run the same analysis scripts for the simulated and empirical data, we randomly generated 23 subjects using the above procedure, so as to be able to run the same standard group analysis using the ADAM toolbox on both datasets (Fahrenfort et al., 2018).

### 2.3. Data preprocessing and analyses

Before applying MVPA analyses, slow drifts were either not removed at all (‘raw data’), or removed through either high-pass filtering, regular robust detrending, or trial-masked robust detrending, each of which is described in more detail below. Each of these preprocessing options was then also analyzed with and without baseline correction.

#### 2.3.1. Removing low-frequency drift noise with high-pass filtering

To investigate the effect of drift removal using high-pass filtering, we high-pass filtered the continuous data (both the simulated and the re-referenced empirical time series) according to typical M/EEG preprocessing pipeline settings. We used one-pass non-causal zero-phase Windowed sinc FIR filter by using EEGLAB’s pop_firws() function (Delorme & Makeig, 2004) with a Kaiser window type, a maximum passband deviation of 0.1% and a stopband attenuation of −60 dB (recommended by Widmann et al., 2015). Filter order was set to correspond to 3 cycles of the cut-off frequency (defined as half amplitude, i.e., −6dB), as recommended in Cohen, M.X. (2014, p. 181). We show the decoding of “raw” unfiltered data, as well as the effect of high-pass filters on decoding for four cut-off values: 0.05 Hz, 0.1 Hz, 0.25 Hz and 0.5 Hz (when showing diagonal decoding) or three cut-off values: 0.05 Hz, 0.1 Hz and 0.5 Hz (when showing temporal generalization, to maintain a manageable number of plots).

#### 2.3.2. Removing low-frequency drift noise using regular and trial-masked robust detrending

As an alternative to high-pass filtering, we either applied robust detrending (de Cheveigne & Arzounian, 2018) which we refer to here as *regular robust detrending*, or an extension of robust detrending which we termed *trial-masked robust detrending*. Detrending involves fitting an n^th^ order polynomial to the data and subtracting the fit, thereby detrending the data to remove slow-fluctuating drifts (de Cheveigne & Arzounian, 2018). Because the fit can be sensitive to sudden deviations from the slow trend (“glitches”; muscle, motion or electrode-specific artifacts; but also sharp, transient ERPs or synchronized oscillations such as posterior alpha band), two unwanted side effects of detrending can occur. First, the glitch can impose ringing artifacts, similar to what happens in filtering. Second, if a signal of interest is partially or fully captured by a high-order polynomial, one risks affecting real effects in an attempt to remove artifacts. In *regular robust detrending*, an iterative weighting procedure is used to identify glitches which are recognized as outliers of the polynomial trend, and which are masked out prior to applying the polynomial fit (see de Cheveigne & Arzounian, 2018, for details). However, although sharp transients are masked out, given the weak nature of ERPs it is not guaranteed that this procedure identifies and thus masks out cognitive events of interest. Thus, a regular robust detrending procedure might still negatively affect the quality the data either (1) by fitting the polynomial to weak but real ERPs, thus subtracting out weak but real effects or (2) because partially fitting the polynomial to the peak of an actual ERPs might actually cause the polynomial to slightly deviate from the baseline activity around the ERP, and as such displace information from the peak of the ERP onto the surrounding window once the polynomial is subtracted from the time series, potentially causing displaced and reversed patterns at these time points.

Therefore, we extended the robust detrending method by actively masking parts of the data that are deemed to reflect experimentally relevant (e.g. cognitive) events. The final fit is then done on the masked data, and subtracted from all data (masked and non-masked, for an illustration of the procedure see Figure 3, for an illustration on real data, see Supplementary Figure S1). Although such a mask might in some cases make the fit to the entire time series slightly worse, it logically prevents that any experimentally interesting effect can be captured by the polynomial fit and thus influence the time series in unwanted ways. We termed this method *trial-masked robust detrending*. For both the simulated and real dataset we used a pre-set mask to remove the ERPs occurring in current trial from the trend fitting operation, so including the encoding, retention and recall phases (i.e. we set a mask that runs from *t* = 0 to *t* = 6 seconds as to not include any meaningful perceptual, cognitive and/or motor-related dynamics into the polynomial fit); all other surrounding data were left unmasked.

High-pass filters are usually applied to continuous data with sufficient buffer zones before and after the experimental recording, because a low-frequency cut-off results in long lasting edge artifacts that may enter the task-related data. However, robust detrending of a whole recording session of typically more than an hour can be suboptimal: the non-stationary slow trend may be too complex, requiring a certain high polynomial order that is difficult to select a priori. Although de Cheveigné and Arzounian provide no clear recommendation as to how long data epochs should be for optimal detrending, the examples given in their paper show segments in the range of a few hundreds of seconds. Because we did not know a priori what the length of the drifts were in our experimental data, we did some preliminary testing and determined that segmented into padded epochs of 56 seconds works quite well (i.e. trial-related epochs of 6 seconds with 25 seconds of trial data pre-/post-padded). To be able to include all trials, the continuous data were symmetrically mirror-padded with 25 seconds prior to segmentation. Note however, that the duration of a padded trial during detrending does not directly impinge on the frequency of the drift that can be removed (as is the case for filter lengths), as a polynomial can easily fit onto a small portion of an oscillation.

Similar to varying the cut-off frequency for filtering, we varied the polynomial order for detrending using the orders: 1, 10, 20 and 30. For all polynomial orders higher than 1, the data were first detrended with a 1^st^ order polynomial (i.e. in fact removing a linear trend over the entire epoch) to improve the fit of the higher order polynomial (as recommended by de Cheveigne & Arzounian, 2018), also see Figure 3 for an illustration of the procedure from top to bottom for a 30^th^ order polynomial. Because of the robust, iterative fitting procedure in robust detrending, the first detrending step updates the mask with additional time-channel-specific outliers; this updated mask is then used as a pre-mask for the next detrending step. As can be observed in the simulated electrodes in Figure 3, the fit is not necessarily perfect (middle panel) and the drift is not perfectly removed (bottom panel). Detrending is not guaranteed to produce perfect fits, as noise can occur in many frequency spectra that are not necessarily always captured by a polynomial of a given order. For this reason, it might be advantageous to try out different filter orders during drift removal. However, by the analytic logic of the mask procedure, the ERP (the signal) that occurs in the current trial cannot affect the fit. Therefore, any remaining effect on MVPA can be regarded as imperfect noise removal, which by the same logic is evenly distributed across trials and conditions.

Robust detrending was done with the Noise Tools toolbox (http://audition.ens.fr/adc/NoiseTools), using the nt_detrend() function. Note that we have added a detrending function to the ADAM toolbox (Fahrenfort et al., 2018) that applies trial-masked robust detrending, allowing one to easily perform a robust detrending and epoching operation on EEG data in EEGLAB format while masking out ‘cognitive’ events, by internally making use of the nt_detrend function. The ADAM function is called adam_detrend_and_epoch(), and takes as inputs continuous EEGLAB data, a specification of the epoch window, the window in which event take place that should be masked out, and some other parameters. Its output can then be used directly for MVPA first level analyses in the ADAM toolbox. The function also produces a plot of the detrending procedure on a trial in the middle of the dataset, using some illustrative channels with strong drifts, as well as a butterfly plot of the ERP data (the average across trials for each electrode) before and after detrending. An example of such a plot can be found in Supplementary Figure S1. See the help of adam_detrend_and_epoch for further details on how to execute the function.

#### 2.3.3. MVPA Analyses

We performed multivariate pattern analyses (MVPA) on both the real and the simulated data, with the use of version 1.11 of the ADAM toolbox (Fahrenfort et al., 2018) – a freely available script-based Matlab analysis package for both backward decoding and forward encoding modeling of M/EEG data. The latest release of the toolbox is available from Github, through http://www.fahrenfort.com/ADAM.htm. A linear discriminant classifier was trained and tested on each time point either using 10-fold cross-validation for both the real and the simulated data. As classification performance metric we used the Area Under the Curve (AUC), in which the curve refers to the Receiver Operating Characteristic (ROC, Hand & Till, 2001).

For the real dataset, the three image categories of faces, houses and letters were used as classes to train the classifier. As features we used all 64 electrodes for the initial analyses, but we also show an analysis in which we pre-selected 9 occipital channels (PO7, PO3, O1, Iz, Oz, POz, PO8, PO4, and O2). This was done to reduce overfitting to irrelevant electrodes, which has been shown to increase classification performance in other visual tasks as well (Fahrenfort, van Leeuwen, Olivers, & Hogendoorn, 2017). Classes comprised the three balanced picture categories, and a linear discriminant classifier (LDA) was used to discriminate the three classes (Fahrenfort et al., 2018). After applying the various preprocessing options (high-pass filtering or detrending), but prior to MVPA, the EEG data were down-sampled to 32 Hz sampling rate using MATLAB’s downsample function (which does not apply an anti-aliasing lowpass filter, to prevent negative temporal effects of lowpass filtering). Another standard pre-processing step is to apply baseline-correction, involving the subtraction of the average activity in a baseline window from the entire trial. Just like high-pass filtering or detrending, this step generally removes unwanted effects of slow noise fluctuations. However, this step may also obfuscate any effects that might be introduced into the baseline window by filtering or detrending. We therefore chose to run MVPA analyses on the data both with and without baseline-correction to be able to compare the two.

Further, we tested the classifier not only on the same time point at which it was trained, but also across all other time points, to inspect effects of the various pre-processing options on temporal generalization. Such across-time decoding generates temporal generalization matrices, which are informative as to whether a pattern of neural activity underlying classification performance is stable, or whether it dynamically changes over time (King & Dehaene, 2014), for example to see whether the activity in the encoding phase is similar (generalizes to) the activity in the retention period. In the context of the current analyses they are also informative with respect to the degree to which patterns are artificially distorted over time. At the group level, subject-specific AUC as a performance measure of multivariate classification was statistically compared against chance for the raw data, as well as for the different cut-offs and polynomial order values using t-tests. For all analyses, we corrected for multiple comparisons using a False Discovery Rate (FDR) correction (q=0.05) on standard t-tests (p<0.05), using a test that is guaranteed to be accurate for any test dependency structure, as described in Benjamini and Yekutieli (2001). This test does not suffer from some of the problems that common cluster-based permutation tests have (Maris & Oostenveld, 2007), as these may have inaccurate onset- and offset boundaries (either in the temporal and/or spatial domain) due to stochastic variation in the noise (Sassenhagen & Draschkow, 2019). Note however, that prior to peer review we had initially analyzed the data using cluster-based permutation and that these analyses showed qualitatively similar results.

For the simulated dataset, the two condition labels assigned to the trials as described in section 2.2.1 served as classes. Other than the number of classes, the analyses were identical between real and simulated data.

## 3. Results

### 3.1. Empirical EEG data

Figure 4 shows classifier performance for the working memory task. We were able to reliably dissociate multivariate patterns of broadband EEG activity across the 64 included channels, during encoding, retention, and the search period for the face, house and letter stimuli. Classification increased transiently during the presentation of the initial target cue, after which it decreased yet remained at above chance levels for up to two seconds during the delay period, before it dropped to near-chance levels. Classifier performance then increased again during presentation of the search display, presumably upon attentionally selecting the target category.

The panels from left to right reveal how decoding performance was affected by different high-pass filter cut-off values (left panel), standard robust detrending orders (middle panel) and trial-masked robust detrending orders (right panel). We also show the effects of these detrending operations both before baseline correction (Figure 4A) and after baseline correction (4B). High-pass filtering and detrending contribute strongly to overall decoding performance during the encoding/retention phase when no baseline correction is applied (the graded effects of increasingly strong cut-offs/polynomial orders in shades of orange when compared to raw decoding in black in Figure 4A), whereas the advantage from filtering or detrending after a baseline-correction is much smaller, at least for the encoding phase (Figure 4B). This is not surprising, given that baseline-correction can be understood as a crude method to achieve the same thing as filtering or detrending: the removal of drifts offsets that occur in a trial. In the case of filtering this happens by attenuating power in the frequency bands in which drift occurs, in the case of detrending this happens by estimating and subtracting out the drift that was estimated by fitting a polynomial, and in baseline-subtraction this happens by subtracting the average (drift-induced) offset of the signal just prior to the presentation of the first target stimulus from the entire trial. Although applying baseline correction has a clear positive effect on decoding performance, it has the disadvantage that its artifact-correcting ability wanes further in the trial, when more time has progressed since the baseline window. This can be observed when comparing the encoding phase and the search phase in 4B, where filtering and detrending have a selective advantage for the strength or extent of above chance decoding performance in the search phase. Such a selective benefit can be particularly problematic in paradigms such as the attentional blink paradigms, when comparing conditions in which different amounts of time have elapsed since baseline correction (short vs long lag, e.g. see Fahrenfort, van Leeuwen, et al., 2017).

Further, note that filtering has small but unmistakable spurious effects on decoding performance, at least for higher cutoffs. For example, FDR corrected significant decoding appears before stimulus onset for a high-pass filter of 0.5 Hz (Figure 4A). Although this cutoff seems to have beneficial effects during the working memory retention period after baseline correction (Figure 4B), we have no way of ascertaining whether this improvement is real. A more likely interpretation is that the baseline correction procedure itself displaces the spurious decoding that was observed prior to stimulus onset in Figure 4A to the remainder of the trial. Normally, the baseline is computed from a window which is assumed to contain no experimentally relevant information. Thus, by subtracting the average activity in that window from the entire time series (separately for every electrode and every trial), one offsets any noise fluctuations prior to stimulus presentation. However, when this window contains a (temporally displaced) pattern of activity that is not due to noise, the inverse of this pattern is transposed once more to the remainder of the trial when applying baseline subtraction, which can now drive decoding accuracy in the rest of the trial. This interpretation seems plausible, as spurious decoding after baseline correction now also emerges prior to the baseline window starting already around −1500 ms. Often, decoding performance is not plotted prior to the baseline window itself, potentially obfuscating such spurious effects. Thus, one may question which decoding effects are real, and which are caused by displacements due to a combination of both filtering and baseline-correction.

A much better grasp of these displacements may be obtained by inspecting the temporal generalization matrices in Figure 5 (without baseline correction) and 6 (with baseline correction). Figure 5A shows temporal generalization for the raw data. Here, the multivariate pattern was significant for the encoding phase (indicated with the number 1 in Figure 5A), but for the raw data did not reach significance after FDR correction for the search phase (number 2 in 5A), nor for generalization from encoding to search and vice versa (numbers 3 and 4 in 5A). High-pass filtering clearly improved this, as both phases (1 and 2) and their generalizations (3 and 4) reach significance after FDR correction for all three cutoffs. Unfortunately however, these improvements come at a cost, as we now also observe spurious FDR-corrected generalization from the encoding and/or search phase to the pre-stimulus window and vice versa for all cutoffs (blue regions left and below the dotted lines), which should logically not be possible given that the pre-stimulus window should not contain information. Thus, this below chance decoding performance indicates that information was displaced from other parts of the trial to the pre-stimulus period. It is also interesting to note that these displacements result in negative decoding performance, i.e. the classifier performed below chance for these temporal generalizations. Plausibly, this is due to the fact that the filtering operation caused displacements to be inverses of the patterns from which they originated (e.g. positive voltages became negative voltages and vice versa), thus explaining a negative decoding performance when generalizing from prestimulus to encoding/search and vice versa.

**Figure 5.**
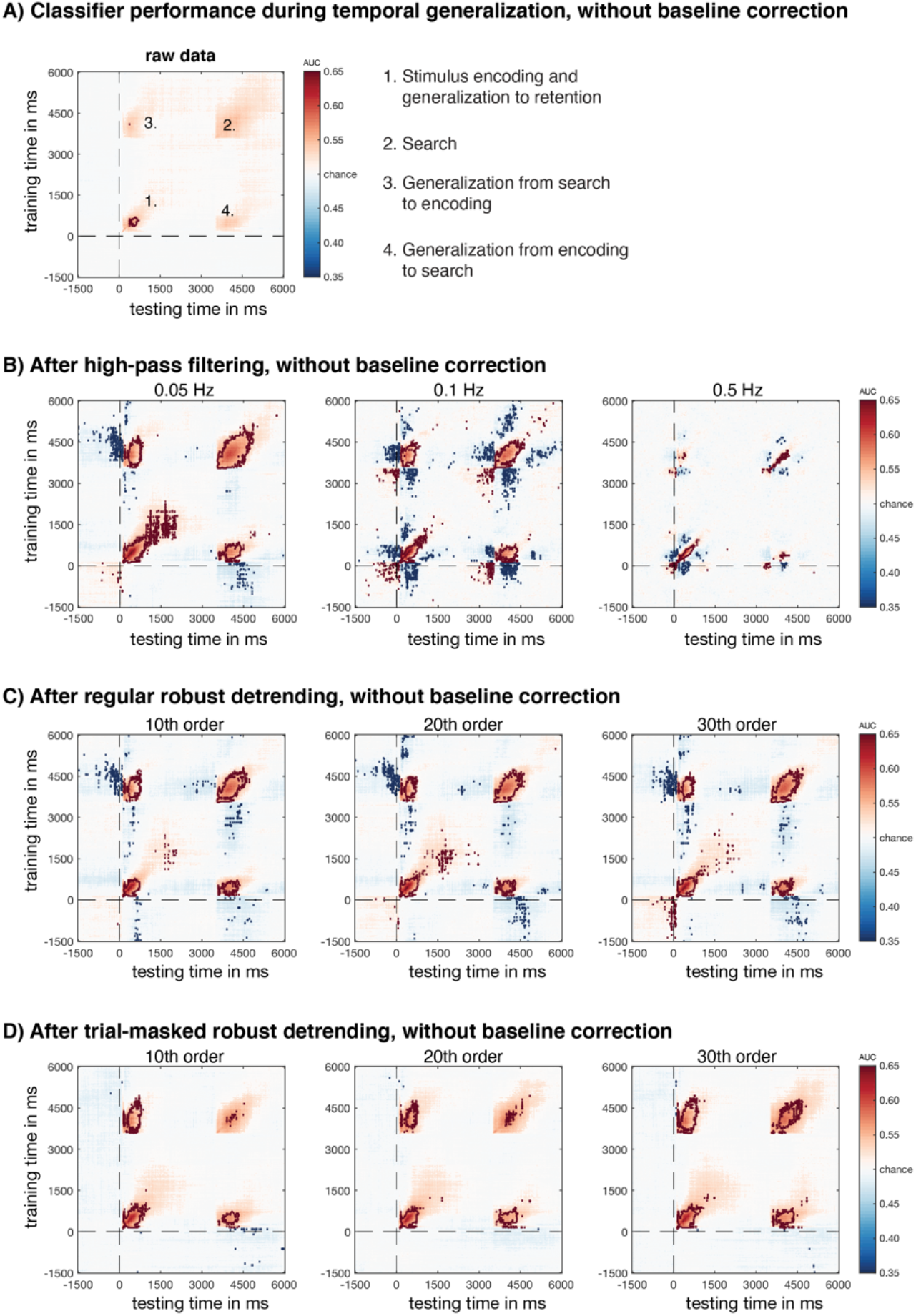
Temporal generalization results for the empirical data without prior baseline-correction. A) Temporal generalization plot for the raw data. Saturated colors are p<0.05 (uncorrected), time points marked or surrounded by dark red (above chance) or dark blue (below chance) contour lines are FDR corrected at q=0.05. The dotted horizontal and vertical line indicate t=0. There should logically be no generalization from the prestimulus window to other points in the trial, so no significant activity left or down from the vertical and horizontal line. The four numbers in the plot indicates various phases in the trial, as explained using the infigure legend. The pattern in the raw data suggests a relatively strong representation during encoding which survives FDR correction, but which does not significantly generalize to the search phase or vice versa (3 and 4) after FDR correction. B) Temporal generalization after high-pass filtering at three different cut-off levels. Note that the encoding phase now significantly generalizes to the search phase and vice versa for all frequency cutoffs. Worryingly though, we now also observe spurious decoding during generalization of the encoding phase to the pre-stimulus window for all frequency cut-offs, even after FDR correction. Blue colors denote below chance decoding during generalization, plausibly due to displaced and reversed patterns caused by filtering, questioning effects observed at other points in the trial as well. C) The same temporal generalization analyses, but now after regular robust detrending at different orders. As for filtering, we observe FDR-corrected significant generalization of the baseline window to the search phase, plausibly caused by pattern reversals during the baseline window due to the detrending operation. D) Trial-masked robust detrending, showing no sign of spurious generalizations, while still obtaining better generalization of the encoding to the search phase and vice versa when compared to raw data.

Further, although Figure 4B seemed to show that no spurious effects occurred for standard robust detrending, we now also observe FDR corrected spurious pre-stimulus effects during temporal generalization of the pre-stimulus window to the search phase in Figure 5C, for all polynomial orders. As for high-pass filtering, these are likely caused by temporal displacements caused by the standard robust detrending operation in similar ways as filtering. The only preprocessing option that did not cause such FDR-corrected spurious decoding accuracies was achieved by trial-masked robust detrending (Figure 5D). In contrast to high-pass filtering and standard robust detrending, the improvements as a result of trial-masked robust detrending occurred without similar increases during baseline periods.

To investigate the effect of such spurious effects in a standard pre-processing pipeline, Figure 6 shows temporal generalization after the same pre-processing steps, but this time after also applying a baseline correction on the basis of the window [−200, 0] ms. Again, it is easy to see how baseline-correction improves decoding performance of raw data when inspecting Figure 6A, both for the encoding and for the search phase (number 1 and 2), as well as for generalization from encoding to search and vice versa (number 3 and 4), all of which now reach statistical significance after FDR correction. Furthermore, the spurious decoding effects that we observed in Figure 5B and 5C as a result of high-pass filtering and regular robust detrending are no longer visible in Figure 6B and 6C. Crucially, the reason for the disappearance of these spurious effects is the fact that the baseline correction procedure itself has moved these spurious pre-stimulus patterns to the rest of the trial, so that they no longer show up as spurious effects in the pre-stimulus window.

**Figure 6.**
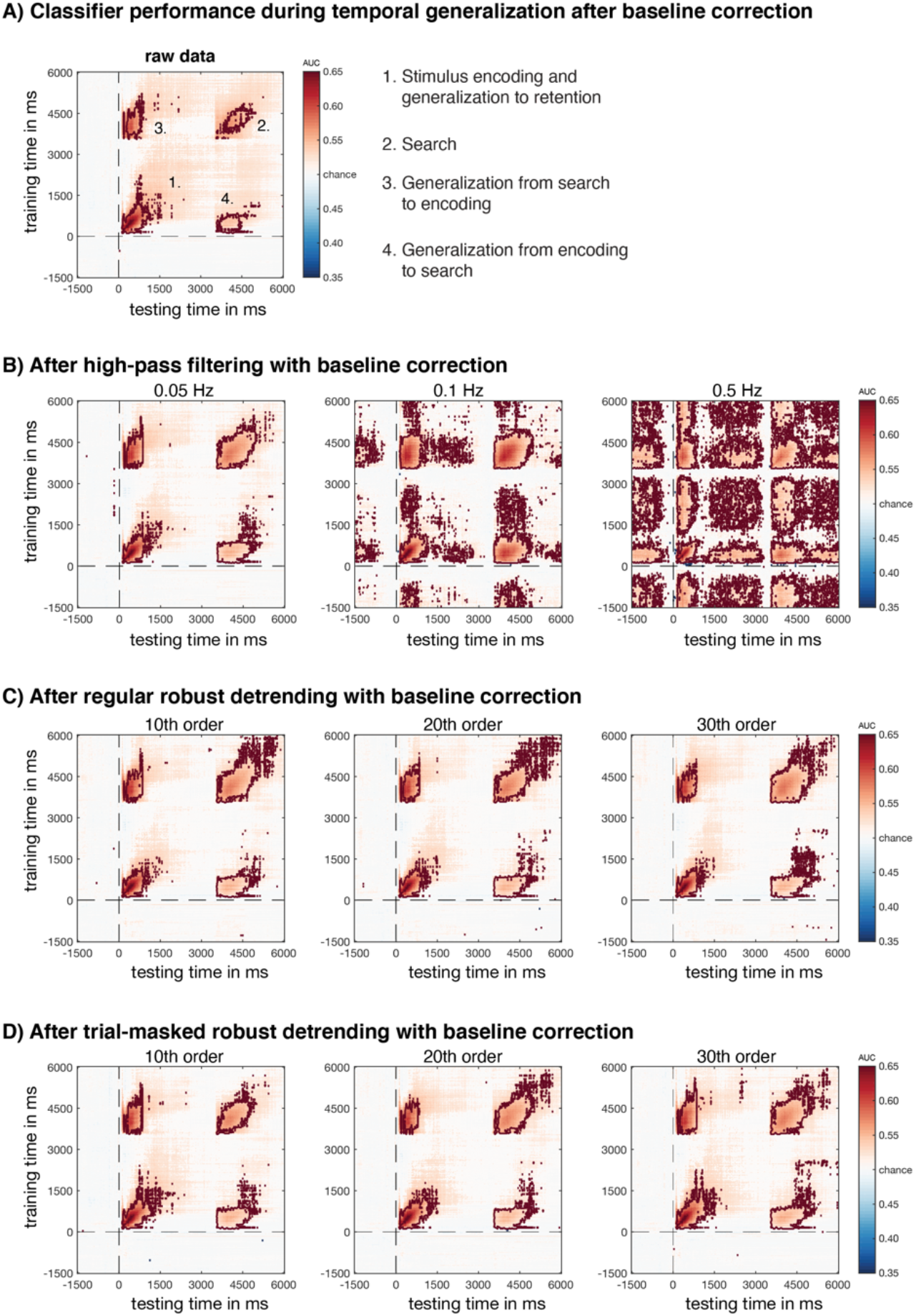
Temporal generalization results for the empirical data with prior baseline-correction. A) Temporal generalization plot for the raw data. Saturated colors are p<0.05 (uncorrected), time points marked or surrounded by dark red contour lines are FDR corrected at q=0.05. Except for an added baseline correction prior to decoding these plots are identical to Figure 5. We now see much better generalization in the raw data from encoding to search and vice versa due to the added baseline-correction. B) Temporal generalization, but now after high-pass filtering at three different cut-off levels. Although the cutoff of 0.05 is seemingly clean, we know from Figure 5 that there were temporal displacements which are now obfuscated by baseline-correction. This becomes very clear when considering the high-pass filter cut-offs of 0.1 and 0.5 Hz, where we now find strong decoding in the part of the pre-stimulus window that was not part of the baseline-window (−1500 to −200 ms), and also in other parts of the trial (retention), plausibly caused by baseline-correction induced temporal offsets of information from the baseline window. C) Temporal generalization after robust detrending and baseline correction. These data seem clean, but from Figure 5 we know that there were offsets in the baseline window even after FDR correction. These offsets are transported back into the rest of the trial due to to baseline correction resulting in potentially unpredictable and unidentifiable spurious improvements in temporal generalization. D) Temporal generalization after trial-masked robust detrending. There are no spurious decoding results and the results seem very similar to decoding of raw data with baseline-correction, barring some seeming minor improvements in the extent of the search phase for 20^th^ and 30^th^ order trial-masked robust detrending.

Thus baseline-correction does not actually resolve the spurious effects that are observed in Figure 5B and 5C, but by its subtraction logic it just moves these spurious effects to the remainder of the trial, making it impossible to disentangle which effects are real and which are spurious. This can be observed when inspecting temporal generalization at high-pass filter cutoffs of 0.1 Hz and 0.5 Hz. These spurious effects of generalization from baseline period to the encoding and search phase now clearly show up as positive decoding in the pre-stimulus window starting at −1500 ms up to −200 ms, both for the 0.1 Hz and for the 0.5 Hz cutoff. This is plausibly what also drives the strong decoding in the retention phase and after the search phase, which can most clearly be seen for the 0.5 Hz cutoff. Further note that baseline correction not just displaces patterns, but also inverts them. For example, a pattern that is described by the vector [−4, 2, 0, −3, 3] becomes [0, 0, 0, 0, 0] after baseline subtraction, but introduces the pattern [4, −2, 0, 3, −3] into the remainder of the trial by virtue of the subtraction logic that is applied. This explains why generalization from the baseline window to the encoding and search phase now turns up as positive decoding for filter cutoffs 0.1 Hz and 0.5 Hz, as the inverted patterns that occurred in the baseline period due to filtering and detrending are inverted once more and end up in the rest of the trial.

Again, it can be easy to miss the spurious nature of such effects if the plotted baseline period is too narrow or if the effect in the pre-stimulus window is constant over time. In either case, baseline correction transports these spurious effects to the rest of the trial without ever having actually observed them. But even when plotting the broader time course, spurious effects in the baseline period may be obfuscated by baseline correction. Indeed, when we inspect the baseline corrected plots after standard robust detrending in Figure 6C, nothing seems wrong with them, even though we know from Figure 5C that there must be spurious effects in the plot, we just have no way of knowing where they are. None of these problems occur for the trial-masked robust detrending plots in Figure 5D and 6D which are free from temporal displacements of events of interest due to the logic of the preprocessing operation, which prevents experimental effects from influencing preprocessing. The fact that the plots in 6C and 6D look similar bring nothing to bear on this fact, Figure 6D is actually the only correct plot from which to draw any conclusions.

One may even question whether there is any advantage to be had from preprocessing relatively clean EEG data at all, as the raw baseline-corrected data in Figure 6A seems to show a similar decoding result as the trial-masked robust detrending data in Figure 6D. However, for subtle effects, it may be beneficial to apply such a preprocessing operation after all. For example, WM representations during retention may be hard to pick due to their low signal to noise ratio. Although there is some debate about where WM representations are maintained in the brain (Christophel, Klink, Spitzer, Roelfsema, & Haynes, 2017), there is strong evidence that visual representations are stored in visual regions during retention (Albers, Kok, Toni, Dijkerman, & De Lange, 2013; Harrison & Tong, 2009; Super, Spekreijse, & Lamme, 2001).Thus to identify such signals (as was the original goal of the data that are presented in the current study), it might be beneficial to apply the decoding operation only to the electrodes on visual areas (occipital). Such a selection might minimize the ability of the classifier to overfit to noise information in electrodes that contain little relevant information. In Figure 7, we show the effect of decoding on only the occipital electrodes (see methods), separately for raw data, 0.1 Hz high-pass filtered data, standard 30^th^ order robust detrending, and 30^th^ order trial-masked robust detrending. Again, we observe the deleterious impact of high-pass filtering, but we also see the advantage of robust detrending over decoding on raw data, especially in the search phase. Further, we observe a clear selective advantage of trial-masked detrending in the temporal generalization from encoding to the retention phase after trial-masked robust detrending, but not after standard robust detrending. Possibly the low signals during retention are subtracted out by including these signals in the polynomial fit during standard robust detrending, but not during trial-masked robust detrending. Highly similar results were obtained for other polynomial orders and cut-off frequencies (see supplementary Figure S2).

**Figure 7.**
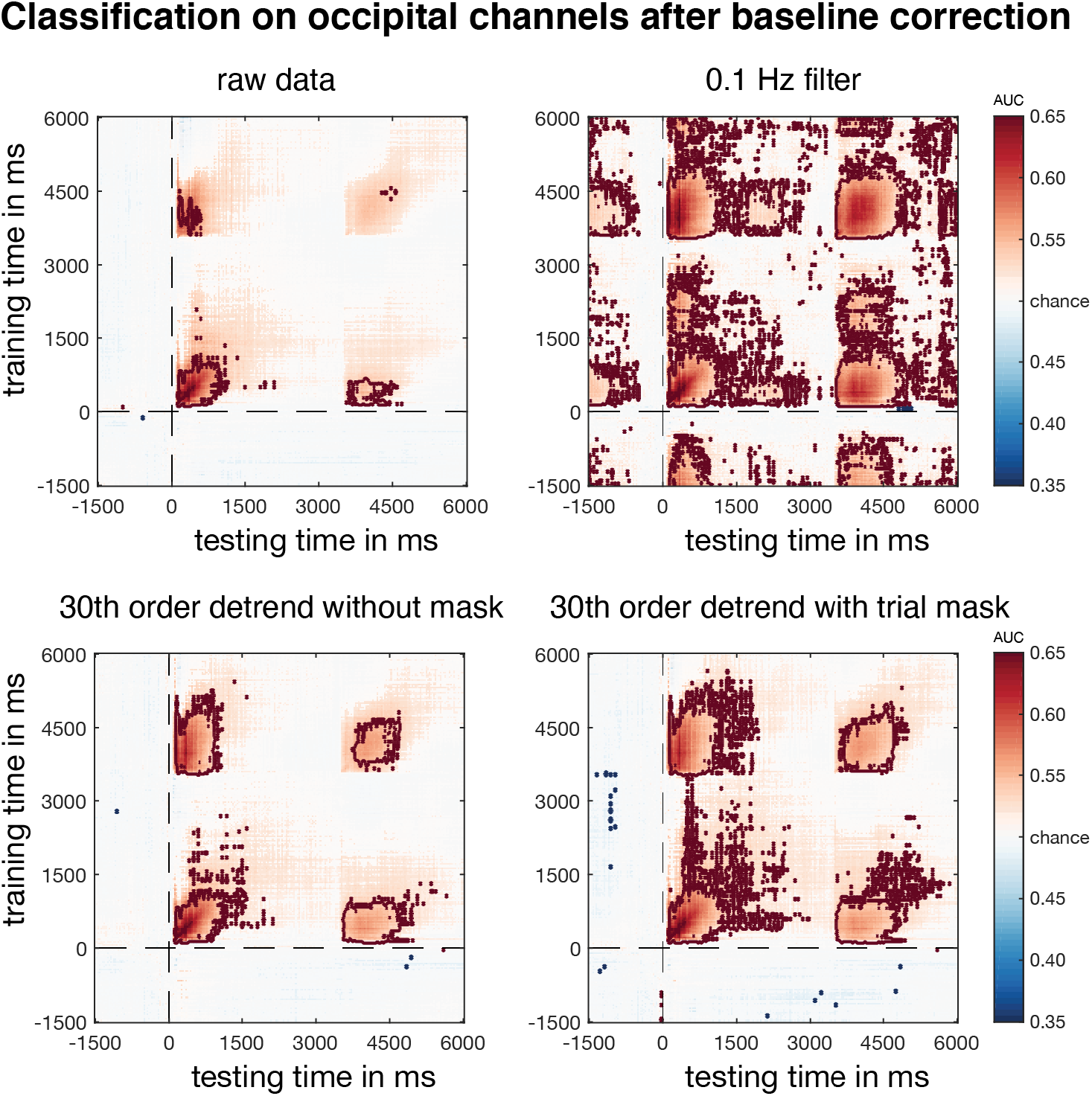
Temporal generalization in baseline-corrected occipital channels under different pre-processing options. The top left panel shows raw data, the top right panel shows 0.1 Hz high-pass filtered data, the bottom left panel shows 30^th^ order detrended data without pre-set trial mask, and the bottom right panel shows 30^th^ order detrended data with pre-set trial mask. Saturated colors are p<0.05 (uncorrected), areas surrounded by dark red contour lines are corrected through an FDR cutoff of q=0.05. The 0.1Hz high-pass filtered data shows clear spurious above chance FDR-corrected decoding performance in the baseline window during temporal generalization in the encoding and search phases due to a combination of high-pass filtering and baseline-correction. Note that 30^th^ order detrending without mask might or might not contain spurious decoding accuracies resulting from similar displacements of information, and in addition might suffer from subtracting out weak information contained in the ERP, while 30^th^ order trial-masked detrending is guaranteed to be clear from such influences, as shown in Figure 5 and 6. Interestingly, 30^th^ order trial-masked detrending with shows better generalization of encoding to retention, plausibly due to the fact that weak signals during retention do not contribute to the fit in trial-masked detrending, so that they cannot be subtracted out.

Summarizing, we find that high-pass filtering can result in clear artifacts in decoding, while contributing little to overall decoding performance. In contrast, trial-masked robust detrending shows no such artifacts, while it does modestly enhance temporal generalization across time when decoding on occipital channels. For subtle cognitive and/or perceptual phenomena, such a small yet significant increase may be very valuable. However, without a ground truth, one may always wonder whether observed improvements are real or spurious. We therefore turned to simulated data, as described next.

### 3.2. Simulated data

Figure 8A illustrates the effect of filtering (left panel) and standard (middle) and trial-masked robust detrending (right panel) on a simulated single trial ERP, for different cut-offs (0.05, 0.1 and 0.5) and orders (1, 10, 30) respectively. This clearly shows attenuated drift for higher level removals, but not without a cost. Filtering with higher cut-off values comes with ringing artifacts surrounding the ERPs, some of which extend up to seconds prior to and after the events (cf. Tanner et al., 2015; Tanner et al., 2016) and can potentially transpose multivariate patterns from one point in time to another point in time. Note that the effect of filtering that we highlight in Figure 8A is mostly caused by an interaction between the filter and the ERP itself, as can be seen in Supplementary Figure S3, where we show the effect of filtering on a noise-free trial (left panel).

**Figure 8.**
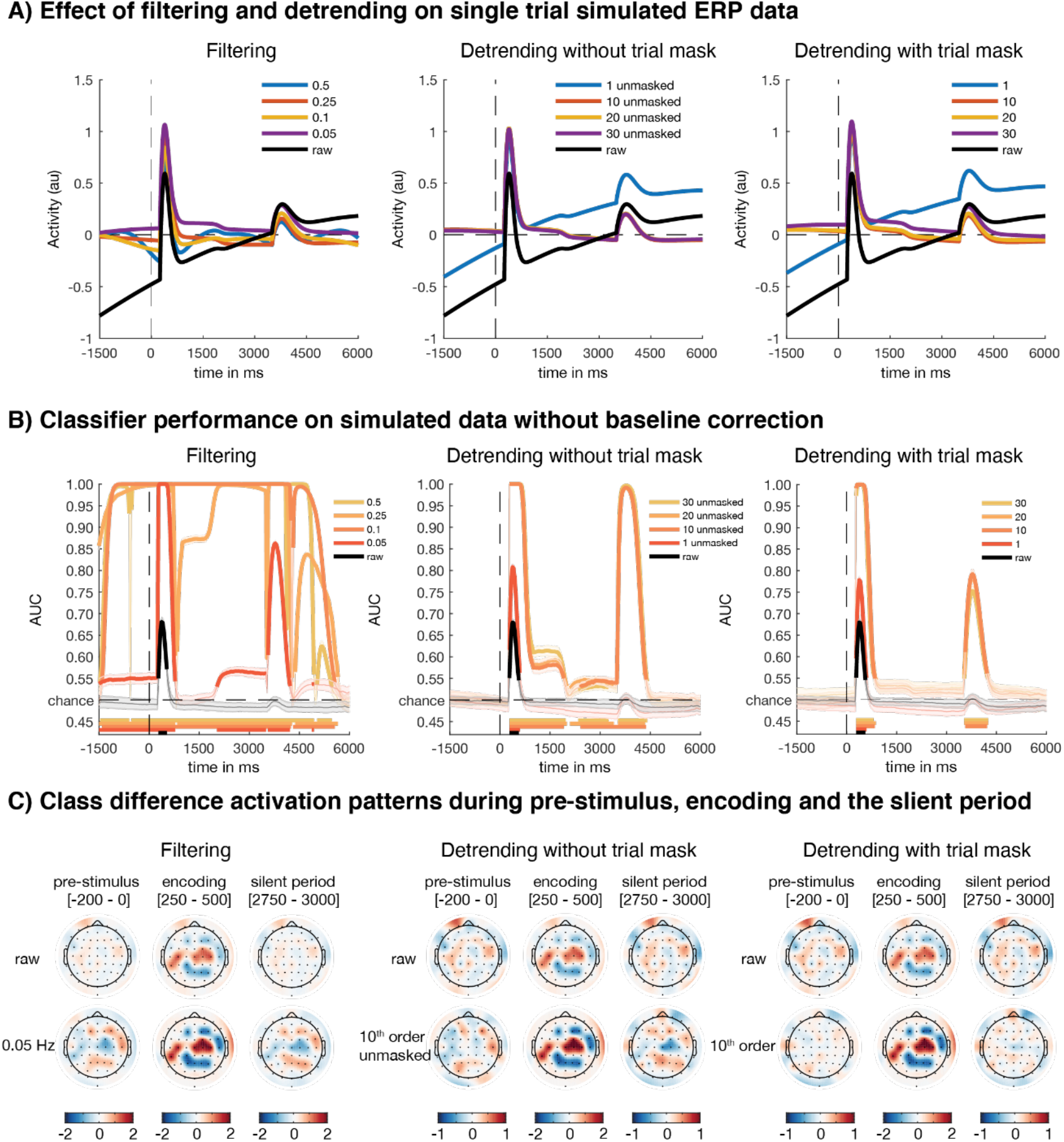
Results for simulated data. A) A single-trial ERP of an illustrative electrode from an illustrative subject for four different high-pass filter cut-offs (left panel) and polynomial orders of regular (middle panel) and trial-masked robust detrending (right panel). Note that no baseline correction was applied, and that the filtering was carried out on the continuous data, while the detrending was carried out on a wider 56.5 second segment, as explained in the methods. B) Decoding performance (AUC) over time for different high-pass filter cut-off frequencies (left panel), regular (middle panel) and trial-masked robust detrending (right panel). Thick colored lines denote reliable difference from chance (p<0.05) after FDR correction (q=0.05). Note the spurious above-chance decoding both prior to the encoding phase (t<0) and in the activity silent interval between encoding and search (between 2.5 and 3.5 s) when no information was present in the simulated data. Similarly, regular robust detrending shows spurious decoding in the activity silent interval between 2.5 and 3.5 s, while trial-masked robust detrending does not. D) Average class-separability maps for different time points in the trial (pre-stimulus, encoding phase, activity silent period) for data that was high-pass filtered at 0.05 Hz (left) and detrended using 10^th^ order polynomials using regular (middle) and trial-masked robust detrending (right). Individual subject patterns were spatially z-scored prior to averaging, color denotes z-value. The encoding phase has a topographical class-separability map that reflects the injected pattern (cf. Figure 2A). Reversed class-related topographical patterns (blue is red and vice versa) can be seen during activity silent time windows when no class-related information was present in the data (t<0 as well as between 2.5 and 3.5 s), both after high-pass filtering data and after regular robust detrending, but not after trial-masked robust detrending.

Although standard robust detrending does not contain such typical filtering artifacts (Figure S3, middle and right panel), it may still result in unforeseen displacements of information, e.g. through an interaction whereby the polynomial fit to a drift is affected by the peak of an ERP. Note here, that transients caused by a regular ERP are typically not masked out by the robust detrending algorithm, especially not when they are embedded in noise that causes the standard deviation to be large (the standard setting detects transient changes of more than three standard deviations from the mean). As a result, standard robust detrending may still adversely affect the data pattern when the polynomial is subtracted from the time series, resulting in the transposal of information present in the ERP to the remainder of the trial. An illustration of this can be found in Figure 8A, where activity in the activity-silent period drops below zero after 30^th^ order standard robust detrending (middle panel) but not after trial-masked robust detrending (right panel)^3^. As explained before, applying a trial-masked robust detrending procedure prevents information from the ERP in the current trial to contribute to the detrending operation. The simulated data we present here are intended to establish to what extent potential effects of different pre-processing operations can be established when the ground truth is known. By using a somewhat caricatural simulated dataset for which the ground truth is known, it is easy to ascertain to what extent the ostensibly spurious FDR-corrected decoding effects we observe in the empirical data are an accidental feature of our empirical dataset, or indeed spurious effects caused by high-pass filtering and/or regular robust detrending.

Figure 8B shows how various high-pass filtering cutoffs (left panel) and polynomial orders for standard robust detrending (middle panel) and trial-masked robust detrending (right panel) affect decoding performance for non-baselined simulated data. Especially filtering has a striking effect on decoding performance, with strong spurious decoding throughout the trial in activity silent windows (t<0 as well as 2.5<t<3.5 s) for all filter cutoffs. Naturally these effects are much stronger than what would be expected in empirical EEG data (cf Figure 4A), but the goal here is merely to show that such spurious decoding effects can potentially result from high-pass filtering. A slightly cleaner picture emerges during standard robust detrending, with decoding performance mostly following the shape of the simulated ERP. However, here too we observe spurious decoding performance in the activity silent window between 2.5 and 3.5 s, drawing into question whether decoding ‘improvements’ in other parts of the trial are real or not. These spurious effects can be observed for all filter orders except the 1^st^ order polynomial (which is a straight line). The only method that shows no spurious effects is trial-masked robust detrending, underpinning the artifact free nature of this method.

When plotting the class-separability maps (topographical maps of the forward-transformed classifier weights, equivalent to the topographic univariate difference between the conditions, Haufe et al., 2014) for filtering/detrending (0.05 Hz and 10^th^ order polynomial, Figure 8C), we see that the encoding phase has a topographical class-separability map that looks identical to the injected pattern (cf. Figure 2A), confirming that both the simulation and decoding operation work as expected. Further, we observe that even for these relatively modest settings, this class-separability map was temporally displaced onto the pre-stimulus window and activity silent period (left panel and middle panel), while this did not occur after trial-masked robust detrending (right panel). Importantly, the displaced patterns are inverted (red regions in the encoding phase become blue regions in the pre-stimulus and silent period, while blue regions become red), in line with the inversions that we observed due to filtering and standard robust detrending in the real data. The strength of these displaced patterns is even stronger for other other filter cutoffs (see Supplementary Figure S4).

To investigate the extent to which these effects are comparable to what we observed in the real data, we also plotted the temporal generalization matrices of the simulated data (Figure 9). These show a very similar (albeit caricatured) pattern to what we observed in the temporal generalization plots of the non-baselined empirical data (Figure 5). Most notably, we observe FDR corrected negative decoding in the generalization windows from baseline to the encoding phase and activity silent periods for both the high-pass filtered and regular robust detrended data, showing up as blue ‘bands’ in a hash pattern (#), which can also be observed in the real data. Notably, we also show that such spurious effects are not observed when applying trial-masked detrending.

**Figure 9.**
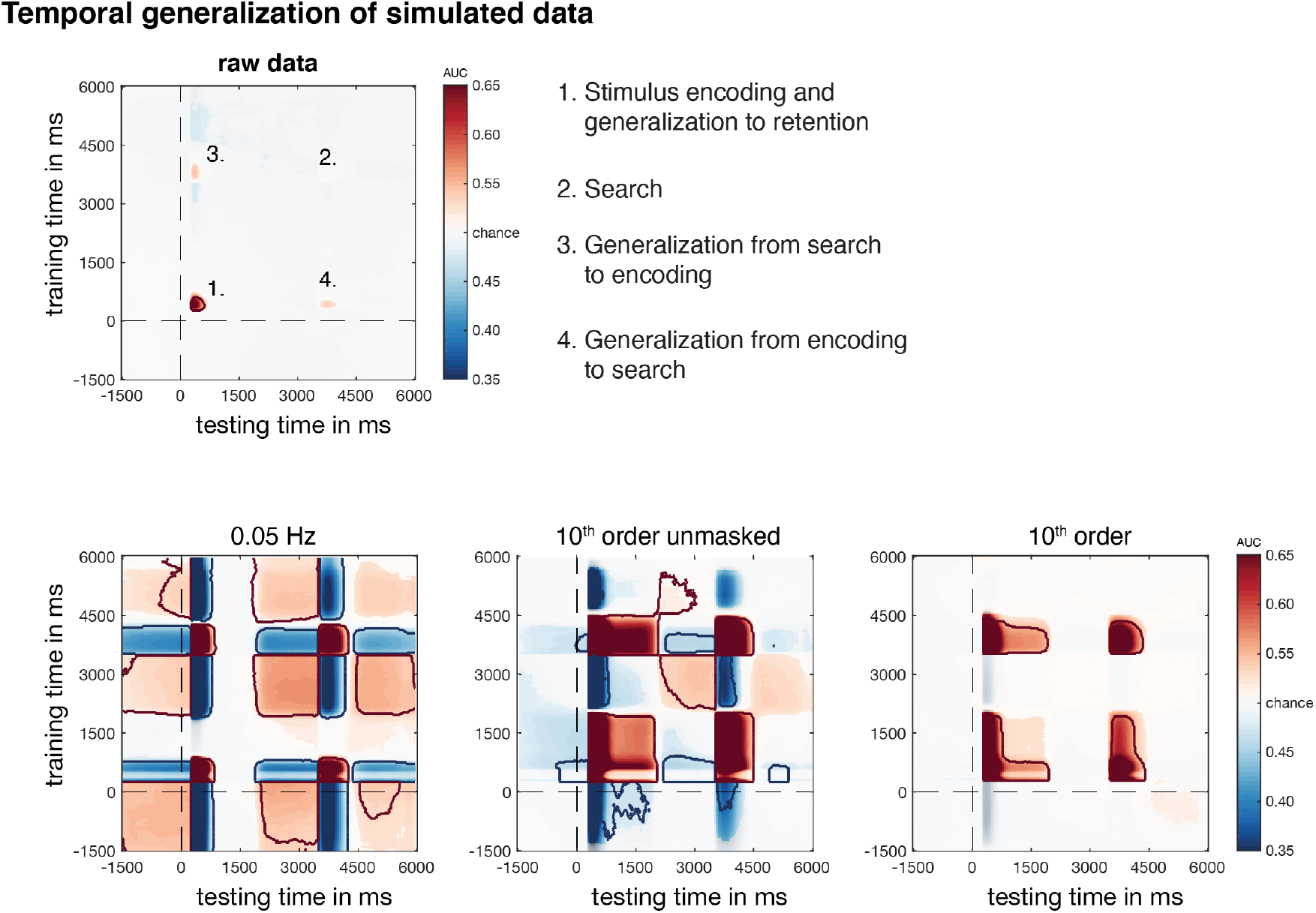
Temporal generalization for the simulated data, without baseline correction. Uncorrected significant p-values are saturated, FDR corrected p-values (q = 0.05) are encircled by dark red contour lines. The raw data (top panel) shows clear decoding of the stimulus encoding phase (number 1), which generalizes to the search phase and vice versa (numbers 3 and 4) although these do not reach statistical significance after FDR correction. The pre-processed data (bottom panel) shows temporal generalization after a modest high-pass filter of 0.5 Hz (left panel), standard robust detrending (middle panel) or trial-masked robust detrending (right panel) with a modest 10^th^ order polynomial. Despite the modest filter and polynomial, we see strong spurious decoding for both high-pass filtering and standard robust detrending but not for trial-masked robust detrending, for which all four phases are now significant after FDR correction, echoing our observations of the empirical EEG data (Figure 5/6). Strikingly, there is strong spurious negative decoding for generalization from activity silent periods (the pre-stimulus window and the window between 2.5 to 3.5 seconds) to the encoding/search phase and vice versa, showing up as a hash-pattern of ‘blue bands’ of negative decoding on the horizontal and vertical, reflecting the presence of reversed multivariate patterns during the activity silent periods, which we indeed observed in the topographic maps of Figure 8C and in supplementary figure S4. The spurious blue hash-pattern can also be found in the temporal generalization matrices of the empirical high-pass filtered and standard robust detrended EEG data in Figure 5, although in the empirical EEG data these blue bands are much weaker and do not reach FDR-corrected statistical significance.

In sum, these simulations show that topographical information on which the classifier relies can inadvertently be transposed onto pre-stimulus and activity silent time windows when applying high-pass filters or regular robust detrending prior to decoding, thus resulting in artificially inflated and extended above or below chance decoding epochs. Trial-masked robust detrending does not suffer from such displacements, as the ERPs are fully masked out. Instead, it comes with few artifacts and improves decoding for components where there is a real underlying signal.

## 4. Discussion

For a long time, high-pass filtering has been a standard step in processing EEG and MEG data, as it has clear benefits when analyzing event-related potentials. Filtering already comes with potential pitfalls for ERP analysis, especially if one is interested in timing/latencies (also see de Cheveigne & Nelken, 2019). Here, we show here that one should also be careful, if not distrustful, when applying any high-pass filtering in service of multivariate pattern classification, because it can easily lead to spurious above-chance decoding effects. We demonstrate these artifacts to clearly emerge for both empirical EEG and simulated data, and from cut-off values as low as 0.05 Hz. From Table 1 it emerges that roughly two-thirds (47/68) of studies have employed a cut-off value higher than that, with 0.1 Hz remaining the most popular value (29 out of the 57 that applied a high-pass filter).

It is important to point out that the enhancement and temporal generalization of decoding performance was particularly salient for time windows where the real underlying signals were actually weak, in particular the “retention phase activity” of our (simulated) working memory task. In working memory experiments, this is an interval during which the memorandum is no longer present, and the classifier is supposed to pick up on purely mnemonic representations. Any enhancement or temporal generalization of such EEG- or MEG-based “mind-reading” capabilities would be very attractive to researchers, and thus extra caution is necessary. We show that above-chance decoding can easily extend to time points where there was actually no signal, not only during the delay period between phases where the simulated signals contained no information, but also during pre-stimulus and in activity silent intervals. Combine this with the fact that the reverse may also happen (i.e. high-pass filtering may actually destroy a real sustained signal, de Cheveigne & Arzounian, 2018), and the risk of drawing false conclusions on the presence or absence of sustained mental representations becomes more than real.

Robust detrending has been advocated as an alternative to high-pass filtering, but we show here that very similar effects can be observed even when applying robust detrending, both in terms of temporal displacements and in terms of subtracting out weak but real information. The cause of the spurious decoding appears to lie in small yet reliable artifacts caused by the interaction between the filter and the ERPs (Acunzo et al., 2012; Kappenman & Luck, 2010; Tanner et al., 2015; Widmann et al., 2015). We propose similar interactions can take place between a polynomial fit and ERPs, even when the fits use an algorithm that is robust to sudden transients and glitches. It has previously been reasoned that standard robust detrending might not suffer from the displacements that are observed in high-pass filtering because either (1) the robust detrending procedure masks out strong transients (from ERPs or otherwise) that might otherwise affect the detrending procedure or (2) ERPs are too weak to affect the polynomial fit, and thus do not affect the detrending procedure. Here we show that this reasoning does not hold up to scrutiny. Even when ERPs contain strong transients as in our simulated data, there may still be temporal windows with low amplitude that are strong enough to affect the polynomial fit despite not being identified as transients, thus either partially displacing ERPs, partially subtracting ERPs out, or both. Although the robust detrending algorithm does allow one to set a threshold parameter that determines what counts as a transient (the default is a deviation of more than three standard deviations from the mean), without setting a pre-trial mask there is no guarantee that the robust detrending procedure precludes potential influences of experimental events on the detrending procedure.

Depending on the nature of the ERPs and filter/detrending settings, artifacts may have quite diffuse effects, extending for seconds prior to and after the relevant events. Although these issues have been pointed out before in the context of ERP analyses, the problem becomes even more salient when performing decoding analyses, as classifiers may be sensitive enough to pick up on subtle, distributed patterns of displaced information that might only be noticed in averaged univariate ERPs when a study has sufficiently high power and one knows where to look. The effects of filtering/detrending on ERPs and on subsequent decoding becomes especially apparent when considering what may be seen as “baseline shifts”, which are then often corrected for in standard ERP analyses. While these might look to be quite subtle, such shifts can actually lead to strong above-chance decoding prior to stimulus onset or when inspecting generalizations from baseline to other time periods in the trial.

It is equally important to point out that baseline correction does not help here, but rather makes things worse, since the baseline applies to the average of an idiosyncratically chosen pre-stimulus period and thus may a) obfuscate displacements introduced by the filtering or detrending operation and b) reintroduce them as artificial effects to the rest of the trial, and even to the period prior to the baseline. This can most clearly be seen when comparing Figure 5B/C and 6B/C, which show how pre-stimulus artifacts are either obfuscated by baseline correction and/or result in spurious pre-baseline and post-stimulus decoding. This is a clear warning that relatively subtle artificial effects in ERP studies can actually have very large undesirable effects on decoding.

The reason for these artifacts lies in the nature of filtering and detrending operations. When filtering the data, one assumes that the noise (drifts) that one attempts to remove through the filtering operation occur in a different part of the frequency spectrum than the signals of interest. However, in practice this is often an unwarranted assumption, especially when trials have a long duration (as for example in working memory experiments that have a retention interval, in attentional blink or other Rapid Serial Visual Presentation paradigms). In such cases, the frequency spectrum in which the signal resides may overlap with the frequency range in which the filter operates. As a result, the filter may end up distorting the signal of interest and displacing information to periods where nothing occurred in reality. In the data presented here (trial duration 6 seconds), even a relatively conservative filter of .05 Hz (filtering out information with a period of more than 20 seconds) nevertheless produced such confounds. Similarly, in a detrending operation, a slow drift polynomial fit may latch on to ERP events and end up ever so slightly distorting the shape of the polynomial, in fact shifting information in time when this polynomial is subtracted from the data.

A better way to remove slow trends from the data is *trial-masked robust detrending*, in which robust detrending (de Cheveigne & Arzounian, 2018) is combined with carefully defined masks that remove potentially relevant (cortical) sources of information from the fits. This method is advantageous when low frequency noise contributions that occur in the same frequency spectrum as the signal of interest can still be separated temporally. In such cases the signal of interest cannot affect the denoising operation because it is masked out.

We found that trial-masked robust detrending can lead to reliable improvements in decoding, while avoiding the artifacts that come with high-pass filtering or standard robust detrending. Nevertheless, here too there are choices to make, and pitfalls to avoid. One important drawback is the search space for optimally detrending the results, where polynomial order and data segment length may interact in ways that in turn depend on the spectral content of the noise one wants to remove. We found that a polynomial order of 30 in combination with 25 seconds padding on each side worked quite well, but in empirical data, this may be unpredictable and study- and subject-specific, further complicating choices as to which detrending options to employ. Moreover, the method we propose here masks out important epochs to prevent fitting to relevant events, but relies on additional assumptions as to which time windows are important. For our particular empirical data set, the improvement achieved with trial-masked robust detrending, relative to the raw data, was interesting but modest (Figure 7). This may not outweigh the extra decisions and assumptions. Of course, this depends on the quality of the data and the conclusions one is after.

Our findings may have wider implications beyond those for EEG decoding analyses. First and foremost, although here we focused on both simulated and empirical EEG data, our demonstrations may naturally apply to MEG data too, given its similar time series structure. Although slow-drift is usually much less of a problem in MEG, similar high-pass filtering procedures have been applied (see Table 1). Second, the spurious displacements of information patterns will not only affect MVPA-based decoding of EEG or MEG data, but also analyses using inverted or forward encoding models that rely on the same type of information (e.g. Herbst, Fiedler, & Obleser, 2018). Finally, there may be important implications for fMRI analyses too. Here is where MVPA took off, with numerous studies demonstrating sustained mental representations beyond the initial stimulus presentation. High-pass filtering is a standard step also in preprocessing fMRI data, and although event-related BOLD responses evolve at a much slower scale than typical EEG or MEG responses, the typical high-pass filter cutoffs used are scaled accordingly. Notably, where in EEG or MEG typically combine trials with event structures in the order of about 2 seconds with high-pass cut-off values in the order of 0.1 Hz, in fMRI event structures are typically in the order of 20 seconds, while cut-offs used are in the order of 0.01 Hz. Interestingly, after pointing out disadvantages of high-pass filtering in fMRI time series (unrelated to decoding), Kay et al. (2008) similarly proposed detrending through polynomial regressors as a solution.

We also note that the decision on whether and how to apply high-pass filtering adds to a list of other design and data processing factors that may all affect decoding results, including transformation into source space, dimensionality reduction, subsampling, aggregating signals across time, artifact rejection, trial averaging, specific classifier selection, and the specific cross-validation design used (Grootswagers, Wardle, & Carlson, 2017). Most notably within the current context, Grootswagers et al. argued for caution when applying *low*-pass filtering (see also Vanrullen, 2011). With too low cut-off values, low-pass filtering too can cause significant decoding to emerge when in fact no signal exists in the original data. Here, we consciously down-sampled our data without applying a low-pass anti-aliasing filter to prevent such issues.

Further, although the general principle we show here is likely to hold for different filter types, we have only explored the impact of a FIR filter with a Kaiser window here. Other options, such as the common 4th order Butterworth filter (Tanner et al., 2015), may produce slightly different results. In choosing only one particular filter, we have also not considered fundamental differences between filter types. Causal filters for example (such as online filters) only take samples from the past and the present into consideration. Naturally, these can never lead to displacement of information backward in time as observed here, although they can still lead to displacement forward in time. Acausal filters on the other hand (such as the offline filter we used here), incorporate information from the future and the past. These types of filters are particularly popular when filtering EEG, because they are able in principle to filter the data without changing the underlying phase of the signal (filters that combine forward and backward filtering are also called zero-phase filters). However, as we have seen here, the promise not to affect the phase of the signal can come at a significant cost, which is that the causal chain of events that the EEG signal attempts to capture can be compromised. How problematic various filter types are in the context of MVPA remains a question for future research.

In conclusion, filtering or detrending of neural time series data may be problematic in more than one respect, but here we show that it becomes particularly troublesome in the advent of modern decoding methods, as it can create widespread displacement of information onto time points where no information was present. This does not have to be problematic in cases where one is not interested in precise timing information and simply wants to assess experimental differences between trials regardless of their timing and/or if one is interested in run-of-the-mill robust average ERP effects. However, this is highly problematic in cases where one wants to investigate more subtle multivariate effects and how they generalize over time. We also show that while trial-masked robust detrending provides a potential solution, no detrending at all may often be good enough. Some labs for which timing information is crucial do not apply filters because of this reason (e.g. Blom, Feuerriegel, Johnson, Bode, & Hogendoorn, 2020). Based on our current findings we therefore recommend extreme caution with regards to high-pass filtering EEG and MEG time series data for MVPA purposes, in particular when timing/onsets/offsets of effects are important, when using slow paradigms such as found in working memory tasks, and when looking at temporal generalization (where spurious results were very pronounced in our empirical dataset). More specifically, we recommend the following steps:

1. Assess the general data quality (unspecific to condition differences). If the quality is good, consider not doing any form of drift removal at all – whether through high-pass filtering or other methods (Luck, 2005). As our own results show, when the data is good baseline correction is often sufficient, so that decoding is likely to work just fine without removing slow trends.
2. This might not be sufficient when the relevant signal extends over longer periods of time. In working memory tasks for example, the retention interval is relatively long, and therefore easily affected by slow drifts. Further, baseline correction is inappropriate when conditions are compared from different time periods since baseline (such as long and short lag trials in the attentional blink), because the amount of time that has elapsed since baseline strongly affects decoding accuracy. In such cases, one might consider trial-masked robust detrending, in which robust detrending (de Cheveigne & Arzounian, 2018) is combined with a pre-set mask so that the ERP and other potentially cognitive events do not contribute to the polynomial fit. Trial-masked detrending precludes the risk that the detrending operation is affected by relevant signals. This also decreases the risk of throwing out real effects. Using this method, we found a modest improvement in decoding accuracy compared to decoding the raw data, in particular when looking at temporal generalization. Still, this method requires one to be aware of several parameters that may affect the results.
3. If there are good reasons to dismiss steps 1 and 2 despite an interest in timing, and one still prefers standard high-pass filtering, then we recommend to systematically explore the cut-off parameter space to assess when spurious enhancement of decoding starts to emerge, and pick a cut-off value well below that (see also Tanner et al., 2015; Tanner et al., 2016). An appropriate double check would be to inspect the temporal generalization matrix of the decoding results, without prior baseline-correction. If strong generalizations occur to or from the baseline window, the filter cutoff should be reconsidered. Given that we found artifacts emerging with cutoff values as low as 0.05 Hz, our choice would be in the range of 0.01 and lower, but this may be different for different event structures and spacing, as there may also be interactions with the intertrial interval (a topic that we chose not to explore in the current study). But even under conservative filter settings, one should be cautious not to overinterpret the precise timing of decoding onsets and offsets when using any kind of filter.

## SUPPLEMENTARY FIGURES

**Figure S1.**
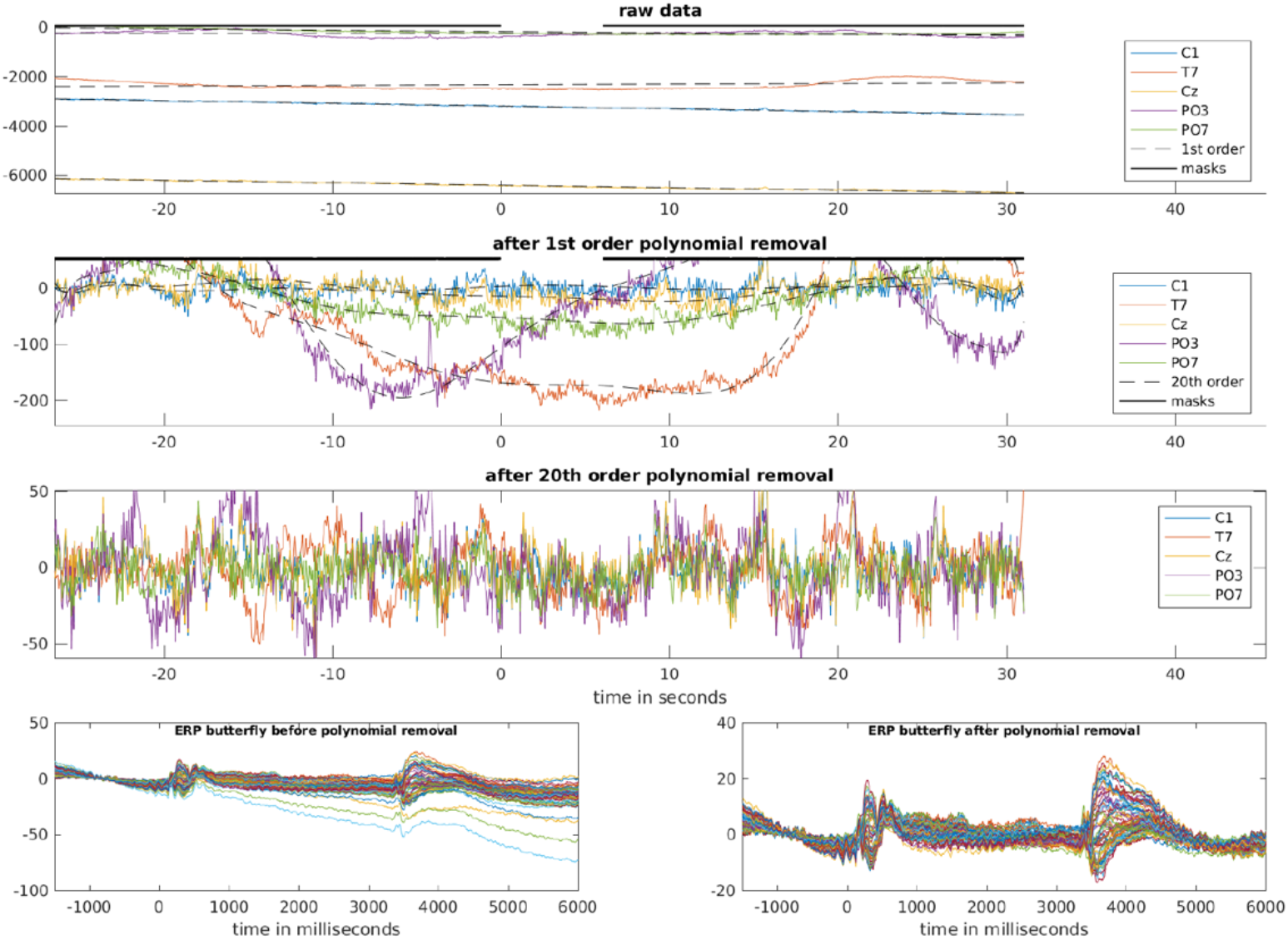
Example figure of a representative subject of the empirical data, generated by the function adam_detrend_and_epoch. The MATLAB function adam_detrend_and_epoch that is included with the ADAM toolbox (Fahrenfort et al., 2018) is able to detrend and epoch continuous data in EEGLAB format. For every subject, it produces a plot like the above. The top three panels show the results of the detrending operation for a trial in the middle of the dataset, for five electrodes that show the largest deviation from center. These three panels are analogous to the panels that are shown in Figure 3 of the main manuscript, but now for empirical data. The bottom two panels show a butterfly plot of the ERPs, i.e. the average ERP of every channel plotted in a different color, both before trial-masked robust detrending (left panel) and after trial-masked robust detrending (right panel).

**Figure S2.**
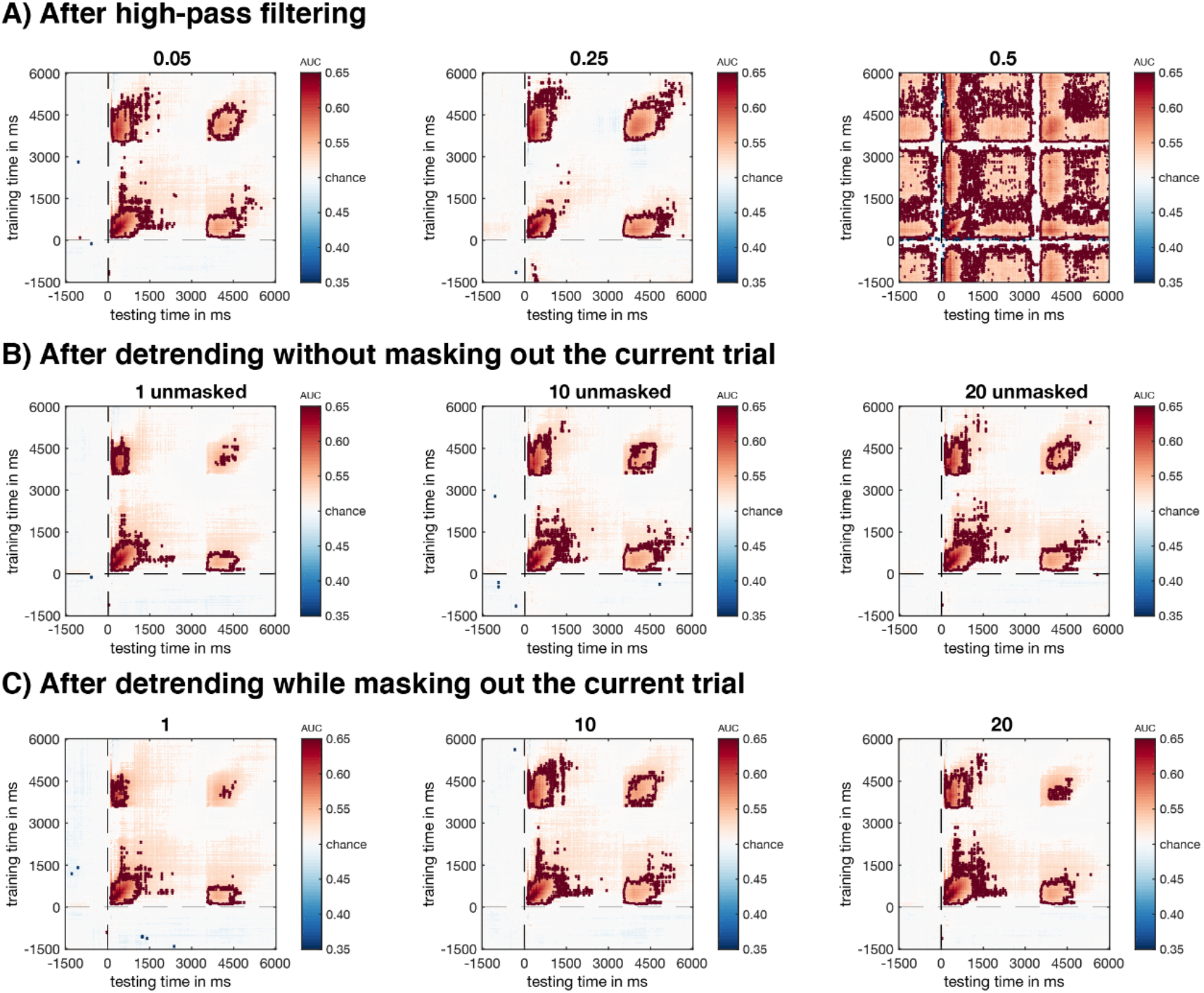
Temporal generalization in baseline-corrected occipital channels under different pre-processing options. Plot shows the same analysis as in Figure 7, but now for different pre-processing options (0.05 Hz, 0.25 Hz and 0.5 Hz high-pass filtered cutoff, as well as order 1, 10 and 20 robust detrended data). The top panels show high-pass filtered data, the middle panels show standard robust detrended data, the bottom panels show trial-masked robust detrended data. Saturated colors are p<0.05 (uncorrected), areas surrounded by dark red contour lines are corrected through an FDR cutoff of q=0.05.

**Figure S3.**
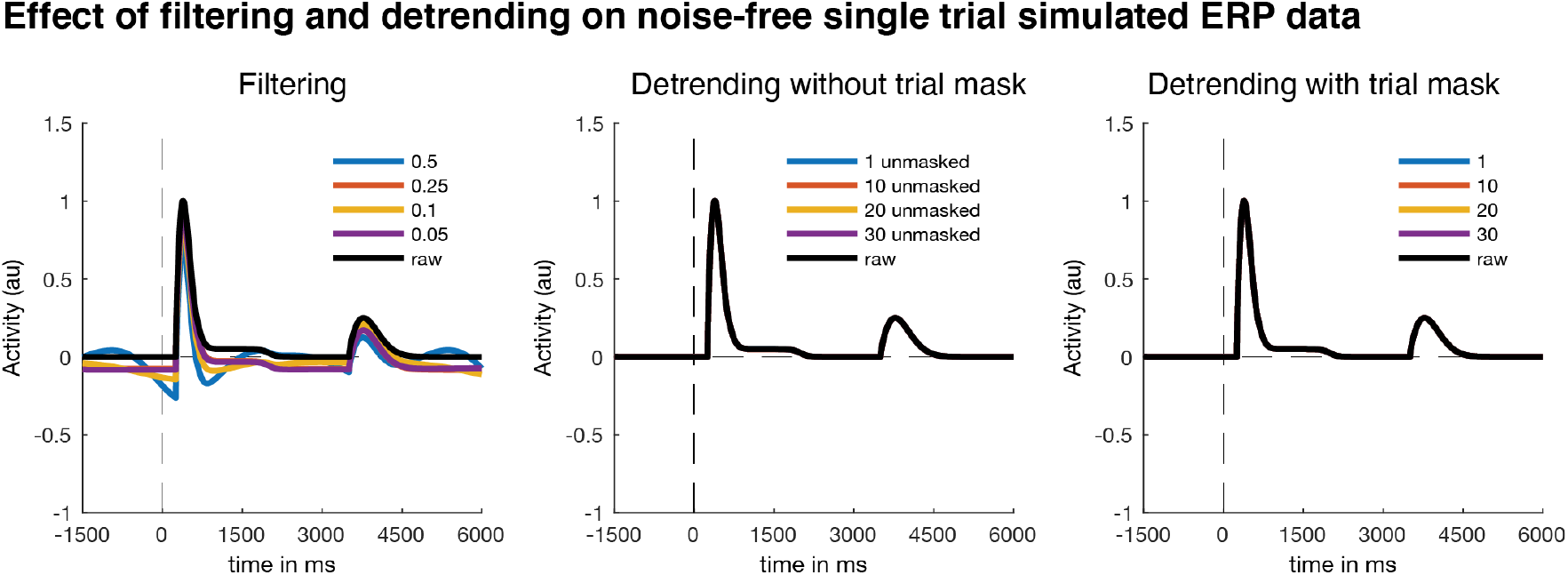
The effect of filtering and robust detrending on a noise free trial (cf Figure 8A). A noise-free single-trial ERP after applying four different high-pass filter cut-offs (left panel) and polynomial orders of regular (middle panel) and trial-masked robust detrending (right panel). Note that filtering was carried out on the continuous data, while the detrending was carried out on a wider 56.5 second segment, as explained in the methods. Further note that here, standard robust detrending (middle panel) seems to give an identical result to trial-masked robust detrending (right panel). This is plausibly caused by the fact that no noise is present in these simulated data, such that even the regular robust detrending algorithm is able to fully and automatically mask out the ERP segments during the detrending procedure, whereas this would be less effective when the ERPs are embedded in noise (as is the case in empirical data).

**Figure S4.**
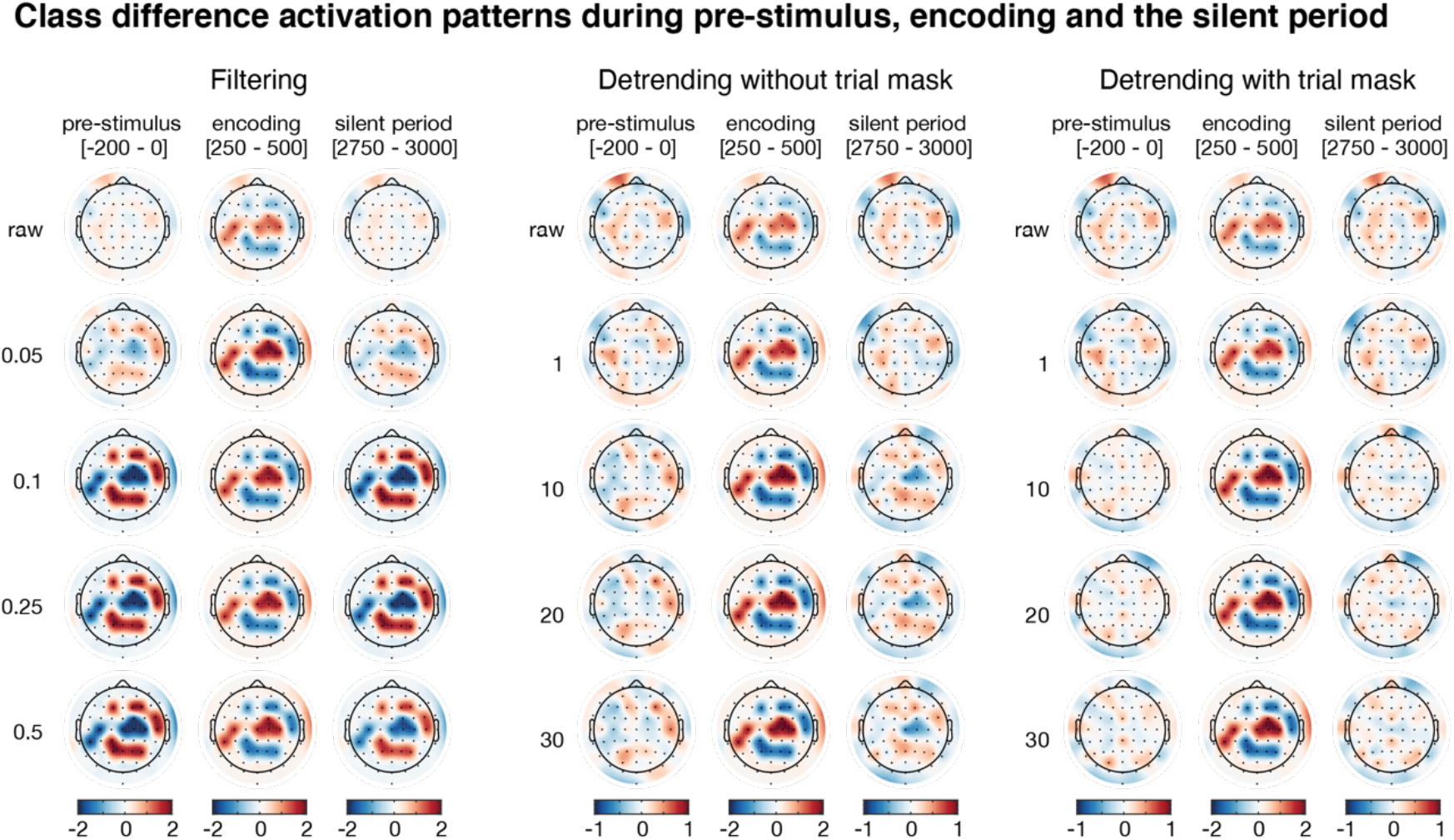
*Average class-separability maps for different time points in the trial (pre-stimulus, encoding phase, activity silent period)* for raw data, data that was high-pass filtered at 0.05 Hz, 0.1 Hz, 0.25 Hz and 05. Hz (left) and detrended using 1^st^, 10^th^, 20^th^ and 30^th^ order polynomials using regular (middle) and trial-masked robust detrending (right). Individual subject patterns were spatially z-scored prior to averaging, color denotes z-value.

## Footnotes

1 Ringing effects are rippling artifacts near sharp edges as a result of filtering out high-frequency information

2 Note that Table 1 is only intended to illustrate the wide-ranging use of filters in EEG/MEG, and not to suggest that anything is necessarily wrong with any particular study. For example, different studies may use different filter types: online (causal) or offline (either causal or acausal), Finite Impulse Response (FIR) or Infinite Impulse Response (IIR), different filter lengths and so forth, and each of these filter types may have different effects on the data that do not necessarily have to be problematic in the scientific context in which they are applied. Here we investigate only one particular common type of high-pass filter in one particular experimental context to assess its influence on MVPA of EEG. We return to this issue in the discussion.

3 Note that we cannot be completely sure of the origin of differences between Figure 8A (middle panel) and Figure 8A (right panel), as they might either be caused by stronger sensitivity to noise outside the trialmask (in the case of trial-masked robust detrending) or by stronger sensitivity to the ERPs themselves (in the case of regular robust detrending), or both. The stronger variability between different orders observed in the case of trial-masked robust detrending (right panel) as compared to standard robust detrending (middle panel) is plausibly caused by a differential sensitivity to noise across outside the trial-mask across orders for trial-masked vs standard robust detrending, but when comparing a differential effect of trial-masked vs standard robust detrending for any given order, this assessment is harder to make.

